# Super compensatory substitutions restore protein function and confer mutational robustness

**DOI:** 10.1101/2025.01.11.631697

**Authors:** Wenhong Jiang, Hengchuang Yin, Saad S. Khan, Yiparemu Abuduhaibaier, Lucas B. Carey

## Abstract

Compensatory mutations can mitigate the fitness costs of deleterious mutations, yet their distribution, structural contexts, and evolutionary roles remain poorly characterized. Here, we systematically analyzed a public deep mutational scanning (DMS) dataset comprising ∼400,000 amino acid variants of imidazole glycerol-phosphate dehydratase (IGPD), with substitutions spanning the entire length of the protein. We found that roughly one-tenth of nonfunctional missense variants can regain function through additional substitutions, predominantly at solvent exposed residues with weak local contacts. Notably, we identified a distinct class of super compensatory substitutions that enhance fitness across diverse genetic backgrounds and buffer mutational effects, resulting in genotypes that occupy locally flatter regions of the fitness landscape. Comparative analyses of additional public DMS datasets across diverse proteins and assays revealed that super compensators recur across different natural sequence–function maps, indicating a broadly represented mode of compensatory effects that sustains mutational persistence and contributes to protein evolutionary robustness.

**Significance:** Compensatory mutations allow deleterious mutations to persist and even fix in populations, shaping protein evolution. Yet, comprehensive analyses of their prevalence and mechanisms have been lacking. Through a systematic analysis of large-scale deep mutational scanning (DMS) datasets, we identified a class of super compensatory substitutions that enhance fitness across diverse genetic backgrounds. Our results reveal a previously unrecognized mode of compensatory effects that promote mutational robustness in evolving proteins.

## Introduction

Most random amino acid mutations destabilize proteins and impair their function (Zeldovich et al. 2007; Faure and Koonin 2015). Nevertheless, deleterious mutations can accumulate and persist within populations through multiple mechanisms that offset fitness costs. Epistatic interactions can mask fitness defects in specific genetic contexts; a mutation that is highly deleterious in one genetic background can become nearly neutral in the presence of another mutation, permitting the accumulation of otherwise deleterious variants (Bershtein et al. 2006; Morton et al. 1956; Lang et al. 2013; Storz 2018), fitting with the view that intraprotein epistasis and folding constraints alter which substitutions are tolerated during evolution (Goldstein and Pollock 2017; Pollock et al. 2012). Proteins can also exhibit mutational robustness, maintaining structural stability and function despite acquiring destabilizing mutations (Bloom et al. 2007; Lind et al. 2016). At the organismal level, allelic redundancy can buffer fitness defects when deleterious mutations affect only a subset of gene copies (Pasek et al. 2006; Gu et al. 2003). Environmental pressures, such as antibiotic exposure, can render otherwise deleterious mutations selectively advantageous, promoting their persistence in populations (Maisnier-Patin and Andersson 2004).

For deleterious nucleotide substitutions, fitness can sometimes be restored through direct reversion to the original sequence. However, when multiple mutations accumulate, or when selective pressures favor retention of the deleterious mutations, additional compensatory mutations provide an alternative route to fitness recovery. These mutations can restore organismal fitness to near wild-type (WT) or even WT levels (Fig. 1 *A*), a phenomenon documented across diverse biological systems (Storz 2018; Maisnier-Patin and Andersson 2004; Bloom et al. 2010; Tomala et al. 2019). Notably, compensatory mutations can occur at higher frequencies than direct reversions (Poon and Chao 2005), particularly following antibiotic-induced population bottlenecks (Meftahi et al. 2016; Comas et al. 2012; Handel et al. 2006; Liu et al. 2018).

**Fig. 1.**
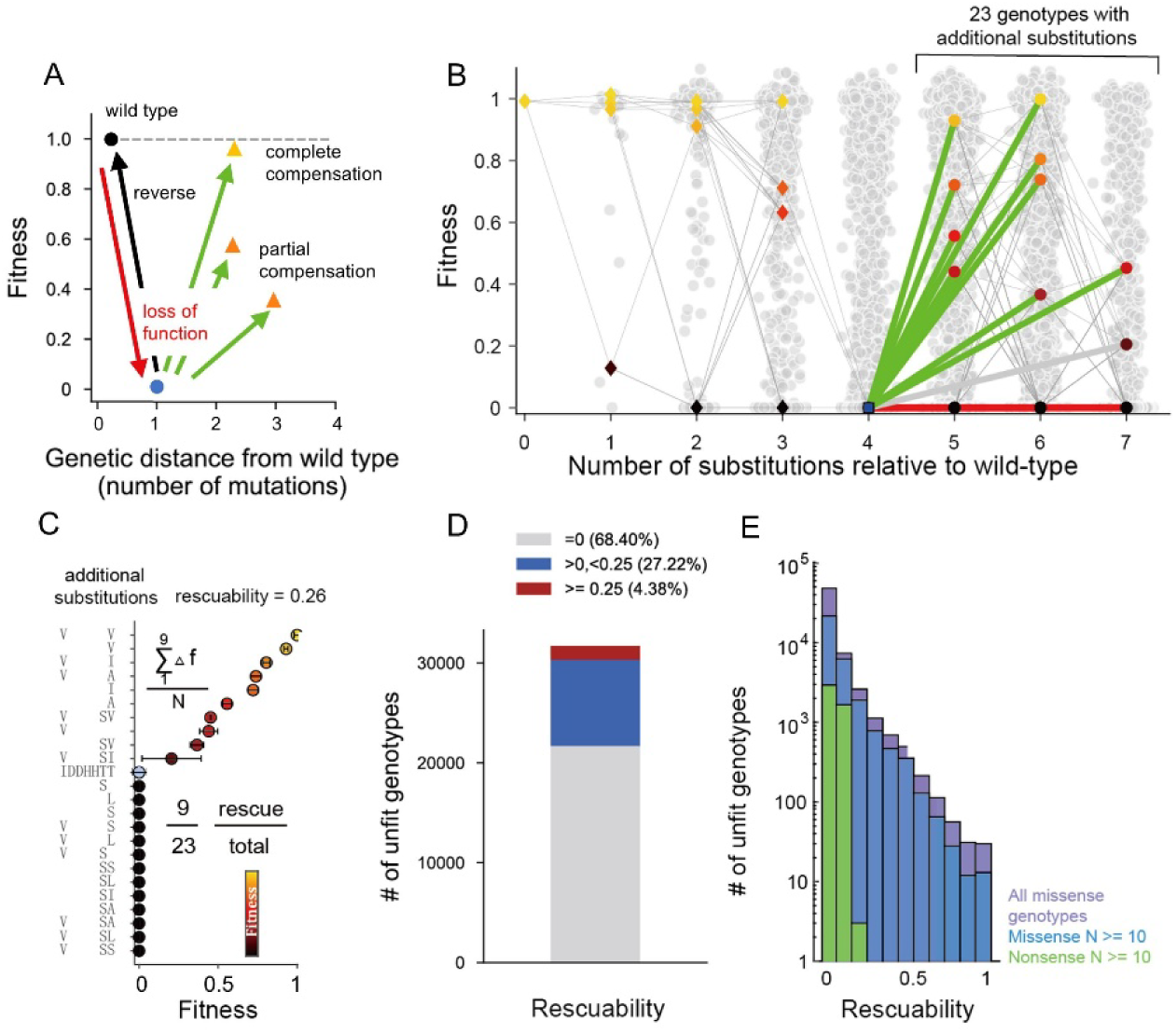
Widespread compensatory substitutions in a deep mutational scanning library. (A) Schematic demonstrating modes of rescue of loss-of-function genotypes via compensatory substitutions. (B) An example of compensatory substitutions in which a genotype with the fitness of zero (blue) is four substitutions away from WT and for which we measured the fitness of 23 genotypes with additional substitutions. The green edges connect to rescuing genotypes, and the red edges connect to non-rescuing genotypes. (C) Fitness data from panel (B) illustrating computations for rescuability. The light blue dot represents a rescuable unfit genotype, whereas the black dots represent genotypes that cannot be rescued. Error bars denote the standard deviation in the estimation of fitness. (D) The distribution of rescuability of all unfit genotypes. (E) Comparison of rescuability distributions among different genotype classes, N denotes the number of synonymous variants used in fitness estimation for the same amino acid genotype. Purple, all missense genotypes; blue, missense genotypes carrying ≥ 10 synonymous variants; green, nonsense genotypes (premature stop codons) serving as a low-rescuability control.

Intragenic compensatory mutations broadly operate through two mechanistic modes: global structural stabilization and pairwise interaction repair. Due to their effect on protein stability, global compensatory mutations have weak spatial constraints relative to destabilizing mutations, provided they contribute to the same folded energetic unit. This spatial flexibility enables broad-spectrum functional rescue across multiple mutational positions (Liu et al. 2018). In contrast, pairwise compensatory mechanisms typically involve residues in three-dimensional contact, forming active-site interfaces, α-helix packing cores, or allosteric communication pathways (Ose et al. 2025). As a result, local compensation is often highly context specific (Ivankov et al. 2014) and can even become destabilizing in the absence of the partnered deleterious mutations (Poon and Chao 2005).

Previous investigations of compensatory evolution have largely relied on small-scale experiments examining the effects of limited sets of genetic variants on individual macromolecules, including proteins (Maisnier-Patin and Andersson 2004; Bloom et al. 2010; Hanson et al. 1993; Mateu and Fersht 1999), mRNA (Stephan and Kirby 1993; Kirby et al. 1995), and tRNA (Domingo et al. 2018). These studies demonstrated that compensatory mutations can restore fitness, but also reported cases where recovery appeared to primarily rely on direct reversion (Filteau et al. 2015; Jakubowska and Korona 2009). Because only a small fraction of sequence space (< 100 genotypes per study) was explored, findings varied substantially across studies (Tomala et al. 2019; Poon and Chao 2006; Ferrer-Costa et al. 2007). This limited sampling has impeded the identification of robust predictors of compensatory potential and obscured broader principles governing the compensation of deleterious mutations.

To address this gap, we analyzed a public large-scale deep mutational scanning (DMS) dataset comprising ∼400,000 amino acid variants of the yeast *HIS3* gene, which encodes the histidine biosynthesis enzyme imidazoleglycerol-phosphate dehydratase (IGPD/His3p) (Pokusaeva et al. 2019). This dataset provides near-complete mutational coverage (>95% of residues) and includes ∼30,000 variants that impair growth in histidine-deficient media. At a scale more than three orders of magnitude larger than previous studies, this dataset enables a systematic exploration of the protein-level principles governing compensatory evolution. Our analysis shows that compensatory potential can be predicted, in part, from protein structural features. Most notably, we identify a distinct class of super compensator substitutions that enhance fitness across diverse genetic backgrounds and buffer the fitness effects of mutations, resulting in genotypes occupying a locally flatter region. The prevalence of these super compensators is further supported by analyses of several additional public DMS datasets. Together, these findings position compensatory substitutions, particularly super compensators, as significant contributors to the shaping of evolutionary trajectories and maintenance of genetic diversity in populations. In this sense, our study provides an experimental perspective on how specific amino-acid states can reshape local sequence context and thereby modulate the accessibility of subsequent evolutionary change (Goldstein and Pollock 2017; Pollock et al. 2012).

## Results

### A DMS Dataset to Identify Intragenic Compensation

To systematically examine the full spectrum of compensatory mutations across an entire protein, we analyzed a large-scale deep mutational scanning (DMS) library of the *Saccharomyces cerevisiae HIS3* gene (Pokusaeva et al. 2019). The *HIS3* coding sequence was divided into twelve 20–30 residue segments, each subjected to localized mutagenesis. Fitness of all variants was quantified through pooled growth assays and normalized such that genotypes carrying premature stop codons were assigned a fitness of zero and wild-type genotypes a value of one (Materials and Methods). After filtering the data for quality (Materials and Methods), the final dataset comprised 372,867 amino acid variant genotypes. Notably, many of these variant combinations correspond to naturally occurring alleles observed across 21 extant yeast species (Fig. S1*A, B*). Among them, 31,701 (8.5%) genotypes displayed a relative fitness of zero and served as the foundation for subsequent analyses (Fig. S1*C*).

In our data, we find that unfit (fitness = 0) genotypes can be rescued by additional substitutions (Fig. 1 *B*). To quantify the extent of rescue, we defined rescuability as the average fitness gained by unfit genotypes upon acquiring one or more additional substitutions (Fig. 1 *C*; Materials and Methods). Using this metric, we find that only a small subset of unfit genotypes exhibits appreciable rescuability (Fig. 1 *D*).

To control for errors in fitness estimation, we used nonsense genotypes containing an internal stop codon as a negative control. Functional rescue of such genotypes is implausible, and indeed we observed little to no rescue among them in our data. Using their rescuability distribution, we estimated a false discovery rate of 5% for rescuability values ≥ 0.02 and 1% for values ≥ 0.04 (Fig. 1 *E*). Based on these calibrations, we classified an unfit genotype as rescuable when its rescuability was ≥ 0.1, identifying 3,750 (11.8%) such genotypes. Notably, comparable rescuability were obtained across a range of fitness cutoffs for definitions of fit and unfit genotypes (Fig. S1*D*) as well as under alternative procedures for calculating rescuability (Fig. S1*E*).

### Characterization of Structural Features Governing Compensation Potential

As nonfunctional variants span nearly the entire protein (Fig. S1*F*), we used the His3p library to evaluate structural features influencing intragenic compensation. To quantify compensatory potential structurally, we defined position-wise rescuability as the mean rescuability of all unfit genotypes carrying substitutions at a given residue (Fig. 2 *A*). Position-wise rescuability varied markedly across residues, with solvent-exposed, core-distal positions, which tend to be weakly evolutionarily conserved, exhibiting significantly higher rescuability (Fig. 2 *B*), consistent with previous observations (Ferrer-Costa et al. 2007).

**Fig. 2.**
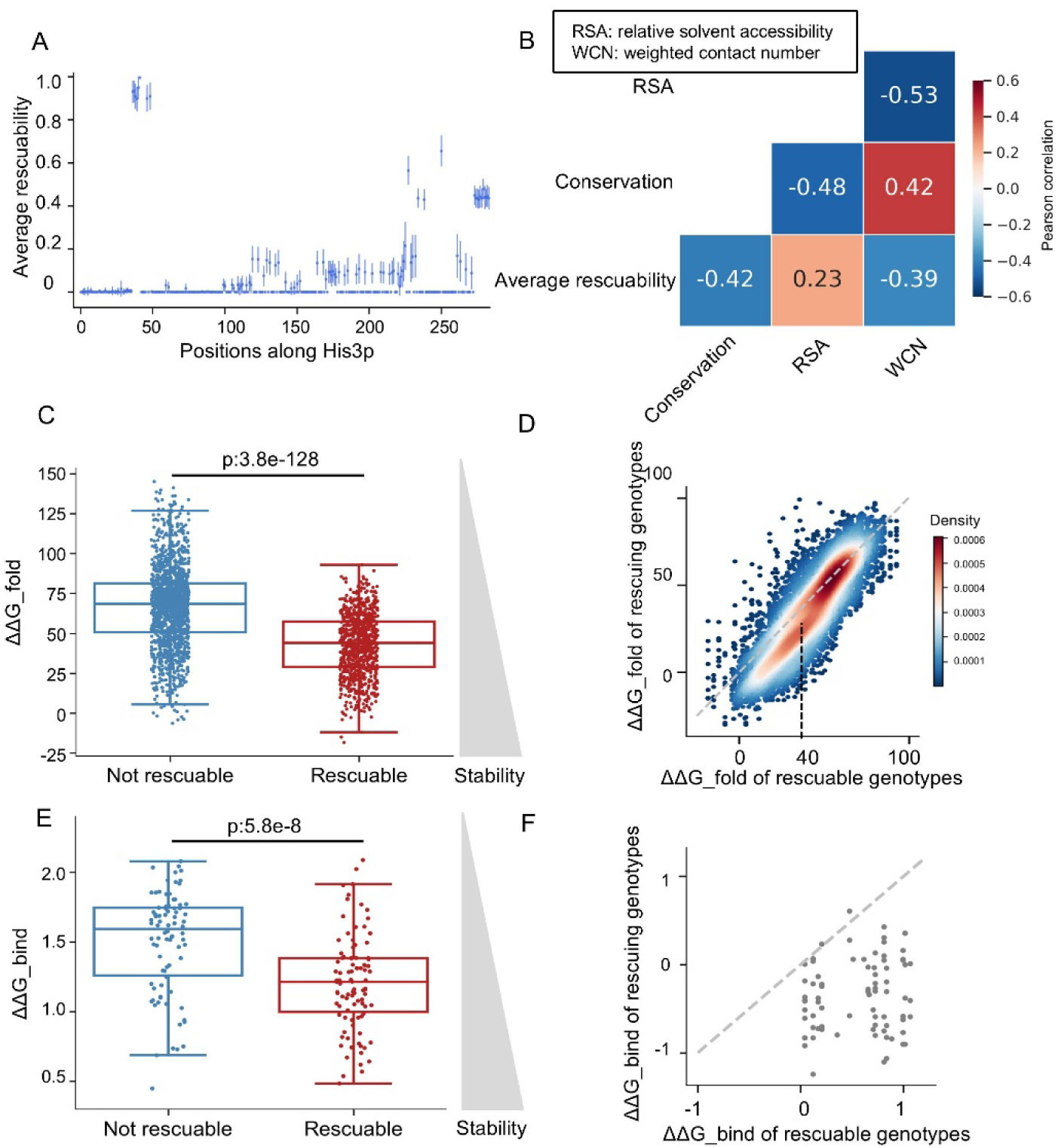
Rescuability correlates with protein structural features. (A) The rescuability of each position in the His3p monomer was calculated as the average rescuability of all genotypes with that position substituted. Error bars represent the standard deviation. (B) Position-wise rescuability correlates with protein structural features and evolutionary conservation. (C) Rescuable genotypes exhibit lower predicted ΔΔG_fold values than non-rescuable ones (one-tailed Wilcoxon rank-sum test). (D) Most rescuing genotypes show lower ΔΔG_fold values than the corresponding rescuable genotypes. (E) Among genotypes with more than 50% of substitutions located in subunit-interface regions, rescuable genotypes display lower ΔΔG_bind values than non-rescuable genotypes (one-tailed Wilcoxon rank-sum test). (F) For genotypes with more than 50% substitutions in subunit interface regions, most rescuing genotypes have smaller ΔΔG_bind than rescuable genotypes.

To investigate the mechanistic relationship between rescuability and structural features, we examined predicted changes in Gibbs free energy change of folding (ΔΔG_fold) for 10,619 representative genotypes. Across all variants, rescuable genotypes exhibited lower ΔΔG_fold than non-rescuable ones, indicating that severe structural destabilization constrains compensatory potential (Fig. 2 *C*; Fig. S2*A*). Moreover, the addition of compensatory mutations partially mitigated destabilization, while restoration to WT stability was uncommon (Fig. 2 *D*). Compensatory stabilization was most pronounced for moderately destabilized variants (ΔΔG_fold < 40), suggesting that structural stabilization contributes significantly, but not exclusively, to intragenic compensation.

Because His3p functions as a homo-24-mer complex (Rawson et al. 2018), we also assessed stability changes at subunit interfaces, where substitutions can influence both monomer folding and inter-subunit binding (Tokuriki and Tawfik 2009; Stein et al. 2019; Casadio et al. 2011). For genotypes with more than 50% of substitution in interface regions, predicted changes in Gibbs free energy change of binding (ΔΔG_bind) mirror the folding analysis: rescuable genotypes had smaller destabilizing shifts, and compensatory substitutions further mitigated losses in binding affinity (Fig. 2 *E* and *F*; Fig. S2*B*). Together, these results suggest that intragenic compensation enhances protein robustness by alleviating destabilizing effects both on monomeric structure and multimeric assembly.

### Super Compensatory Substitutions Increase the Fitness of Diverse Variants

Given the observed structural specificity of rescuability, we next sought to understand modes of compensation across different genetic backgrounds. Specifically, we asked whether compensatory substitutions act in a genotype specific manner or whether some substitutions can restore fitness across a broad range of deleterious genotypes.

To quantify these modes of compensation, we represented intragenic rescue relationships using a directed graph in which nodes correspond to amino-acid states and directed edges encode compensatory effects. An edge from amino acid state A to B indicates that introducing substitution B increases the fitness of genotypes carrying state A. Edge weights represent the fraction of such genotypes rescued beyond a defined fitness threshold (≥ 0.5; Fig. 3 *A-B*). The overall compensatory ability of a substitution was defined as the sum of its incoming edge weights, quantifying the breadth of rescue across heterogeneous genetic contexts. Thus, the network captures how well each substitution can rescue genotypes with distinct existing amino acid states.

**Fig. 3.**
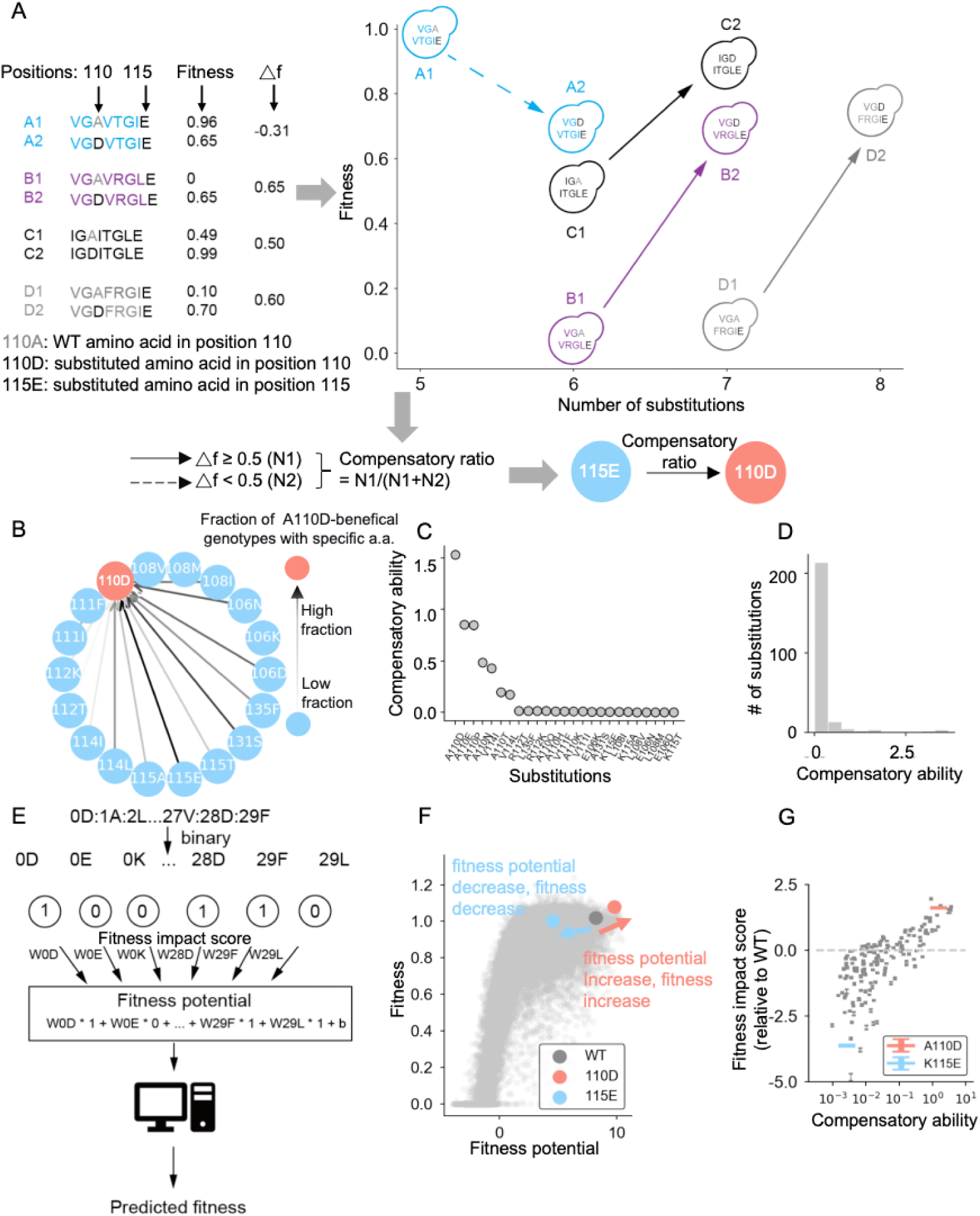
Super compensatory substitutions increase fitness across diverse genetic backgrounds. (A) A schematic figure quantifies the compensatory ability of A110D in the 115E background. For substitution A110D in non-WT state 115E, paired genotypes differing only at position 110 are compared. Fitness differences (Δf) are computed and genotype pairs with Δf ≥ 0.5 are considered compensatory. The resulting compensatory ratio is represented as a directed edge from 115E to 110D. (B) Directed network of compensatory interactions. Nodes represent non-WT amino acid states and directed edges compensatory effects of substitutions. Edge shading indicates fraction of substituted genotypes considered compensatory. (C) Compensatory ability of individual substitutions, defined as the sum of incoming edge weights in (B). Only substitutions from Segment 1 are shown. (D) Distribution of compensatory ability across all substitutions, showing that only a small subset of substitutions exhibits high compensatory capacity. (E) Neural-network model predicting fitness via a latent variable (fitness potential), computed as the sum of amino acid specific fitness impact scores mapped to fitness by a non-linear transformation. (F) An example illustrating modulation of fitness and fitness potential by substitutions: A110D (red) increases fitness potential and fitness, whereas K115E (blue) decreases both. (G) Substitutions with higher compensatory ability tend to exhibit higher fitness impact scores relative to WT.

This analysis revealed a small subset of super compensatory substitutions—specific amino acid changes (e.g., A110D) capable of increasing the fitness of genetically diverse variants (Fig. 3 *C* and *D*; Fig. S3; Fig. S4). The corresponding non-WT amino-acid states resulting from these substitutions (e.g., 110D) are hereafter referred to as super compensators.

To investigate why certain substitutions emerge as supercompensatory, we used a published global-epistasis model that assigns each amino acid state a fitness impact score (Pokusaeva et al., 2019). The model takes one-hot encoded amino-acid states as input, combines them into a one-dimensional additive latent variable termed fitness potential, and then maps this latent value to predicted fitness through a learned nonlinear output layer with 22 units and a sigmoidal transformation (Fig. 3E,F; Fig. S5). In this framework, a genotype’s fitness potential is simply the sum of the fitness impact scores of its constituent amino acid states, providing an interpretable link between state-specific effects and overall fitness. Remarkably, super compensators exhibited substantially higher fitness impact scores than their WT counterparts (Fig. 3 *F* and *G*). These elevated scores enable super compensatory substitutions to increase fitness potential—and thus predicted fitness—across diverse genetic backgrounds, offering an explanation for their broad compensatory action.

### Super Compensators Buffer the Effects of Beneficial Substitutions

To understand how super compensators achieve broad compensation, we examined their influence on the fitness effects of other substitutions. Specifically, we compared the fitness effects of identical substitutions introduced into wild-type (WT) versus super compensator backgrounds. Remarkably, replacing a WT amino acid with a super compensator substantially attenuated the fitness effect of both deleterious and beneficial substitutions at other positions, indicating a strong buffering effect (Fig. 4 *A*).

**Fig. 4.**
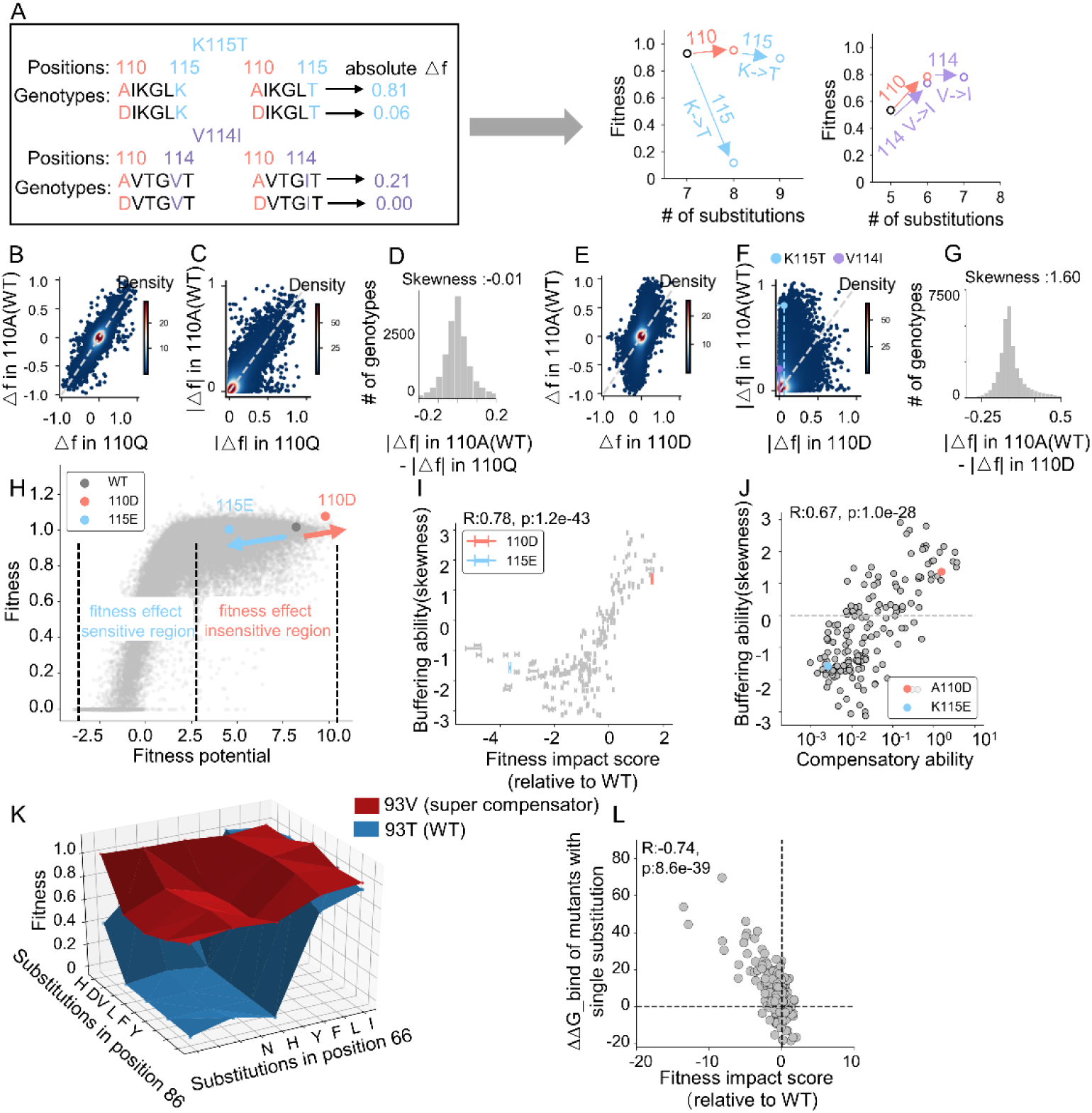
Super compensators buffer the fitness effects of both deleterious and beneficial substitutions. (A) Representative examples showing that super compensator 110D attenuates the fitness effects of both a deleterious (K115T) and beneficial (V114I) substitution. (B-D) A non-super compensator (110Q, x-axis) shows no systemic effect on substitution fitness compared with the WT amino acid state (110A, y-axis). Paired fitness effects were evaluated using relative (B) and absolute (C) fitness difference and skewness of the absolute differences (D). (E-G) A super compensator (110D, x-axis) reduces the fitness effects of substitutions relative to the WT state (110A, y-axis). Comparisons using relative (E) and absolute (F) fitness difference reveal buffering through collapse toward the vertical and produce a positively skewed distribution of fitness effect differences (G). (H) Conceptual model linking buffering to the sigmoidal relationship between fitness potential and predicted fitness. Non-WT amino acid states with positive fitness impact scores (eg., 110D) can shift genotypes towards a high-fitness plateau where fitness is insensitive to additional substitutions. States with negative impact scores (eg., 115E) can shift genotypes off the plateau, increasing sensitivity. (I) Amino acid states with higher fitness impact scores relative to WT exhibit greater buffering capacity, quantified as increased positive skewness of fitness effect differences (Spearman R= 0.78, p-value = 0). (J) Non-WT amino acid states with higher compensatory capacity similarly exhibit stronger buffering, as indicated by increased positive skewness (Spearman R= 0.67, p-value = 0). (K) Example of local landscape smoothing by a super compensator. 93V reduces fitness variability among neighboring variants, effectively flattening the local fitness landscape. (L) Non-WT amino acid states with higher fitness impact scores relative to WT are associated with more stable protein structures, reflected by lower ΔΔG_bind values.

To quantify this buffering effect, we computed the fitness difference (Δf) of each substitution with the WT (Fig. 4 *B*, y-axis) and the non-WT amino acid state (Fig. 4 *B*, x-axis). If substitution effects were comparable in both contexts, data points would distribute symmetrically around the diagonal. Instead, for super compensators such as 110D, the distribution collapsed along the vertical axis (Fig. 4 *E*), demonstrating substantial dampening of substitution effects. We further quantified this behavior by calculating the absolute deviation between the x- and y-axes (Fig. 4 *C*, *F*), and assessing the skewness of this distribution. Substitutions in super compensator backgrounds exhibited both reduced absolute fitness effects and greater skewness relative to non-super compensator contexts (Fig. 4 *D*, *G*), confirming the enhanced buffering capacity of super compensators.

To identify the underlying cause, we revisited the sigmoidal relationship between fitness potential and predicted fitness defined by the neural-network model. Most genotypes in our library reside on the high-fitness plateau, where substantial variation in fitness potential produces only minor change in predicted fitness (Fig. 4 *H*). Beneficial substitutions that push genotypes further along this plateau have reduced sensitivity toward fitness deviations due to additional substitutions, yielding positively skewed fitness-effect distributions. Conversely, deleterious substitutions drive genotypes off the plateau, increasing sensitivity and generating negatively skewed distributions (Fig. 4 *H*). Across all genotypes, we found that amino acid states with higher fitness impact scores than their WT counterparts exhibited higher skewness in the distribution of absolute fitness-effect differences (Fig. 4 *I*). Similarly, substitutions with higher compensatory ability showed correspondingly increased skewness (Fig. 4 *J*). These observations indicate that super compensators buffer both deleterious and beneficial substitution effects by operating within a flattened region of the fitness landscape.

Whereas prior studies have emphasized the ruggedness of protein fitness landscapes, characterized by numerous local peaks and valleys (Kondrashov et al. 2002; Bak et al. 1992; Van Cleve and Weissman 2015; de Visser et al. 2018), our results show that super compensators can make genotypes located in a flatter-than-expected segment of the fitness landscape. The super compensator 93V provides a clear example: in variants containing substitutions at positions 66 and 86, 93V substantially reduces fitness variability among neighboring variants, effectively resulting in genotypes occupying a locally flatter region (Fig. 4*K*).

### Super Compensators Stabilize Inter-residue Interactions

Building on the role of stability restoration in compensation, we next evaluated structural features of super compensatory substitutions. In general, super compensators significantly increased overall protein stability (Fig. 4 *L*; Table S1). For instance, 110D forms a salt bridge with R112, effectively stabilizing a neighboring loop region. In contrast, substitutions exhibiting lower fitness impact scores than the WT, such as I19D, were predicted to disrupt a β-sheet and its adjacent helix (Table S1).

Notably, most super compensators are spatially distant from the catalytic sites (> 8 Å), with none located within 4 Å of the substrate (Fig. S6). This spatial separation supports a mechanism of global stabilization rather than direct modulation of active-site interactions.

### Validating Super Compensators as Robust Buffers of Deleterious Substitutions

To experimentally verify the functional consequences of super compensation, we performed deep mutational scanning to determine whether super compensators confer mutational robustness by buffering deleterious substitutions. We designed the library to contain two fully randomized flanking positions (NNK codons) surrounding a central position encoding either the WT or the super compensator S189A amino acid (Fig. 5 *A*). Under histidine-deficient conditions, variants carrying premature stop codons displayed severely reduced fitness, which returned to WT levels upon histidine supplementation (Fig. 5 *B*), confirming the reliability of our assay.

**Fig. 5.**
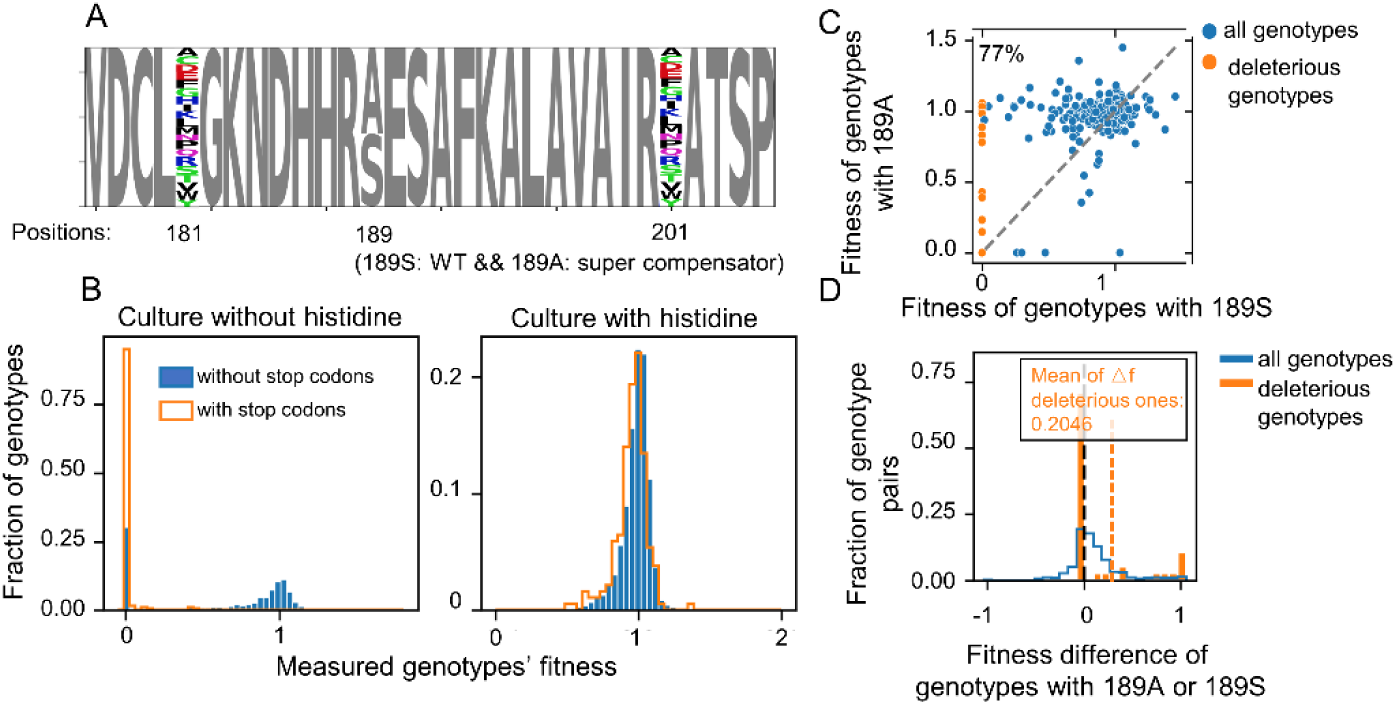
The Super compensator 189A buffers the fitness effects of deleterious substitutions. (A) Amino-acid composition of the constructed DMS library. (B) Genotypes with premature stop codons exhibit similar fitness as those without in the presence of histidine and show reduced fitness in its absence. (C) Super compensators increase the fitness of most genotypes carrying random mutations. (D) The average fitness difference between genotypes with 189A and with 189S is higher than zero (blue), and the fitness difference is more pronounced among deleterious genotypes (orange).

Introduction of the super compensator 189A resulted in a marked fitness increase for 77% of all variants in the library (Fig. 5 *C*). The rescue effect was strongest among genotypes harboring mildly deleterious substitutions (Fig. 5 *D*). Together, these results provide direct experimental evidence that super compensators can increase mutational tolerance by buffering fitness defects caused by otherwise deleterious mutations.

### Super Compensators are Prevalent across Diverse Proteins

To assess whether super compensators represent a general phenomenon beyond the His3p system, we applied the same analytical framework to previously published DMS datasets derived from other randomly mutagenized proteins, assayed using diverse fitness readouts (Notin et al., 2023). Four datasets met criteria: three measured in vitro protein folding stability via cDNA display proteolysis—E3 ubiquitin-protein ligase AMFR, iron-regulated surface determinant protein H (IsdH), and E3 SUMO-protein ligase CBX4 (Tsuboyama et al., 2023)—and one measured fluorescence intensity via FACS (green fluorescent protein 2, GFP2) (Gonzalez Somermeyer et al., 2022).

By applying the same computational pipeline, we identified specific amino acid states with high compensatory ability (Fig. S7A) that consistently buffer the fitness effects of mutations across different proteins (Fig. S7B). These findings suggest that super compensators exist in independent biological systems.

Notably, although fitness was defined differently across datasets—as yeast growth rate in His3p, protein stability in a cell-free system (Tsuboyama et al., 2023), and fluorescence intensity as a proxy for protein folding and activity in GFP2 (Gonzalez Somermeyer et al., 2022)—our results remained robust to these variations. Taken together, these observations support a general model in which super compensators exert their effects, at least in part, by enhancing protein stability.

## Discussion

Our analysis reveals the principles governing intragenic compensation, delineating both the characteristics of rescuable genotypes and compensatory substitutions. We found that the compensatory potential of non-functional variants is strongly correlated with protein structural context: substitutions at solvent-exposed, weakly spatially constrained positions are more easily rescued. This finding clarifies why Tomala et al. (Tomala et al. 2019) identified a limited set of compensatory substitutions for temperature-sensitive mutations, which often reside in structurally critical regions, consistent with independent evidence that some deleterious mutations are fundamentally uncompensatable (Jakubowska and Korona 2009). Consistent with Tomala et al.’s results, we observe that deleterious mutations causing modest structural destabilization can be rescued, reinforcing our observation that the destabilizing effect of substitution in folded and conserved positions is stronger than those in the solvent-exposed positions (Tokuriki and Tawfik 2009; Greene et al. 2001). Beyond structural context, we also demonstrate that amino acid physicochemical properties can serve as predictors for fitness. Specifically, when the AAindex values—a set of quantitative descriptors of amino acid attributes—in mutant variants approximates those of the WT (Kawashima and Kanehisa 2000), fitness recovery is observed (Fig. S8). That is to say, compensation can occur when mutations bring the AAindex values closer to those of the WT.

Focusing on the properties of compensatory substitutions themselves, we identified a distinct class of super compensatory substitutions capable of increasing fitness of diverse genetic backgrounds. This phenomenon parallels the concept of cryptic mutations (Zheng et al. 2019), which facilitate the accumulation of otherwise deleterious substitutions. Our experimental validation directly demonstrates this capacity: introducing super compensators reduced deleterious effects of mutations, thereby increasing mutational robustness, supporting the idea that the effect of a mutation cannot be fully understood in isolation from the surrounding sequence background (Goldstein and Pollock 2017; Pollock et al. 2012). The existence of super compensators can thus provide an alternative explanation for the mutational robustness of proteins. Previous work by Guo et al (Guo et al. 2004) and György Abrusán et al (Abrusán and Marsh 2016) found mutational robustness to vary across protein secondary structures. This aligns with our observation of increased compensation at solvent exposed positions and indicates that structure context shapes both the emergence and effectiveness of compensatory mechanisms.

The evolutionary implications of super compensators may extend beyond laboratory assays. If super compensators can increase the fitness of diverse variants, they should leave a detectable signature in natural sequences by enabling the retention of mildly deleterious alleles. Consistent with this expectation, comparative analysis of 335 His3p orthologs revealed that super compensators frequently co-occur with mildly deleterious amino acid states across extant yeast species (Fig. S9A). Moreover, the strength of the super compensator (its fitness impact score) was positively correlated with the severity of its partner deleterious amino acid states (Fig. S9B). These observations provide a plausible explanation for the sequence divergence observed among His3p orthologs and suggest that super compensators may have contributed to natural genetic variation.

These insights have practical implications for protein engineering. The ability of super compensators to confer mutational robustness suggests that predicting and incorporating them could serve as an effective strategy for designing proteins with enhanced stability and expanded functions. Accordingly, prioritizing stabilizing substitutions represents a valuable component for protein design, consistent with prevailing approaches that emphasize predicting stability-increasing mutations (Popov et al. 2018; Borgo and Havranek 2012; Zimmerman et al. 2017; Steipe et al. 1994).

While the His3p DMS dataset is invaluable, its limitations warrant consideration. First, its basis in extant sequence backgrounds may underrepresent severely deleterious substitutions. Second, segmenting the gene precluded the detection of inter-segment compensatory effects. Nevertheless, to our knowledge, the public His3p library remains one of the largest and most comprehensive resources that contains multi-site mutants (Pokusaeva et al., 2019) and the current work combines a newly generated targeted dataset with a fresh re-analysis of this public His3p library to reveal previously unappreciated rules of compensation. Furthermore, we are able to recapitulate our results regarding the stabilizing role of super compensator across other public large-scale DMS datasets, including those based on GFP fluorescence (Gonzalez Somermeyer et al. 2022) and other stability-directed assays (Tsuboyama et al. 2023). This cross-validation underscores the generalizability of our conclusions beyond His3p.

In summary, this work deepens our understanding of how compensatory evolution is constrained by protein structure, reveals how super compensators promote mutational robustness and sequence diversity, and highlights how these principles can inform rational strategies for protein engineering.

## Materials and Methods

### Experimental Design

The experimental methodology and analytical procedures were already fully described in the previous work (Pokusaeva et al. 2019). We briefly summarize the key steps as follows: first, plasmids containing each *HIS3* gene variant encoding IGPD with distinct amino acid substitutions were constructed and transformed into Δ*HIS3 S. cerevisiae*. Mixed populations of yeast strains carrying different *HIS3* variants were then co-cultured in bulk competition assays. Deleterious substitutions in IGPD resulted in slow growth and decline of variants from the population over time, neutral or beneficial substitutions enabled rapid growth and eventual dominance.

Through growth analysis, variants with missense substitutions exhibiting growth rates comparable to or lower than premature stop codon-containing variants were assigned a fitness value of zero. In contrast, WT genotypes and the majority of single-substitution variants demonstrated fitness values of 1. The experimental design included two biological replicates with two technical replicates per biological replicate. Following rigorous quality control, which excluded data with a replicate correlation of < 0.85, a further filter was applied to ensure robust fitness assessment. Segment 9 exhibited low correlation between biological replicates and was therefore excluded from this work (Pokusaeva et al. 2019). Specifically, amino acid genotypes represented by fewer than three synonymous nucleotide variants were discarded. The resulting dataset consisted of genotypes with an average of seven amino acid substitutions from the reference sequence. The fitness of each genotype was assessed by averaging nine synonymous variants. As these synonymous mutants exhibited little to no fitness difference (Pokusaeva et al. 2019), this internal replication provided high confidence.

### Calculation of Rescuability

For each unfit genotype, we identified all genotypes containing the same set of substitutions plus one or more additional substitutions (hereafter referred to as the superset, total count N). Fitness for each amino acid genotype was estimated using synonymous nucleotide variants, yielding a mean fitness (S) and standard deviation (S.D.). The 95% confidence interval for fitness was defined as S ± 1.96×S.D. A superset genotype A was considered to compensate for genotype B when its lower confidence bound exceeded the upper confidence bound of B, i.e., *S_A_*-1.96*S.D._A_* **≥** S*_B_* +1.96*S.D._B_*. For each unfit genotype, we calculated fitness difference (Δf = S_A_ -S_B_) for all compensating superset genotypes (N1 out of N). Rescuability was then defined as the sum of these Δf values normalized by the total superset count :

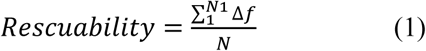

As an independent validation, we used a non-parametric approach using synonymous variants. For each candidate compensating genotype A and unfit genotype B, we compared the distribution of fitness values of their respective synonymous variant sets (G_A_ and G_B_) using a two sample Kolmogorov-Smirnov test (α=0.05). Genotype A was considered compensatory if the fitness distribution of G_A_ was significantly higher than G_B_. Fitness difference (Δf) and rescuability values were then computed analogously to the primary method.

### Calculation of Structural and Evolutionary Metrics

#### WCN Score

The WCN calculation was performed using established methodology from Lin CP et al. (Lin et al. 2008), with the formula:

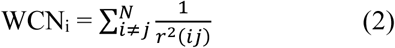

where r_ij_ denotes the distance between the C_α_ geometric center of residues *i* and residues *j* in IGPD (N total residues). The WCN value for each position was calculated by averaging measurements across 24 monomeric structures.

#### RSA Score

The RSA score for each position of IGPD was derived from the DSSP file (generated using IGPD’s PDB file 6EZM) (Carter et al. 2003). The RSA values were normalized using the amino acid-specific surface area reference established by Tien et al. (Tien et al. 2013).

#### △△G Prediction

To predict energetic effects of mutations, we computed ΔΔG for monomer stability using the ddg_monomer protocol (Kellogg et al., 2011), and for protein–protein interface binding using the flex ddG protocol (Barlow et al., 2018).

#### Conservation Score

Conservation scores were derived using sequence entropy analysis of IGPD homologs. Monomeric IGPD sequences from 21 extant species were aligned using T-coffee Expresso (Notredame et al. 2000). For each alignment position X, sequence entropy was calculated as:

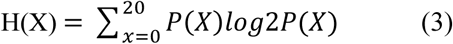

Where P(X) represents the probability of each amino acid at position X, estimated from the multiple sequence alignment. The conservation score for position X is defined as:

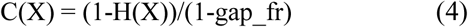

Where gap_fr quantifies the gap occurrence rate at position X.

### Association of Physicochemical Features with Fitness

For each genotype, we quantified physicochemical distance from the wild-type sequence as the difference in AAindex values (AAindex distance) relative to WT across 553 AAindex descriptors. Analyses were performed within mutation sets, defined as genotypes carrying substitutions at the same positions but differing in amino-acid identity. For each mutation set and AAindex descriptor, we computed spearman correlations (ρ) between AAindex distance and corresponding fitness across genotypes. Mutation sets were required to contain at least 82 genotypes, corresponding to the minimum sample size needed to detect modest correlation (ρ=0.3) with α = 0.05 as determined by an a priori power analysis using G*Power 3.1 (Faul et al. 2009). Correlation P values were adjusted for multiple testing across all descriptors and mutation sets using the Benjamini–Hochberg false discovery rate (FDR), correlations with FDR < 0.05 were considered significant. For visualization, non-significant correlations were set to zero.

### Directed-Graph Representation of Compensatory Interactions

For each substitution at position i (S_i_) and each non-WT amino acid state at distinct position j (Nj), we identified pairs of genotypes that only differed at position i: G_1_ = (S_i_, N_j_) and G_2_ = (WT_i_, N_j_). The compensatory ability of S_i_ with respect to N_j_ was quantified as the fraction of such genotype pairs for which Δf = f(G_1_) - f(G_2_) > 0.5. These relationships were represented as directed edges from S_i_ to N_j_ in a network graph, with edge weights corresponding to the fraction of rescued genotype pairs. The visualizations produced for Figure 3*B* and Figure S3*A* were produced using NetworkX (Hagberg et al. 2007) in Python.

### Quantification of Buffering of Substitutions Fitness Effects

To assess how super compensators modulate the magnitude of fitness effect of other substitutions, we considered only substitutions that produced a nonzero fitness effect. The fitness effect of a substitution was defined as the difference in fitness between two genotypes that differ only at a single position (for example, G_1_=00000, G_2_=00100, where 0 denotes WT amino acid and 1 denotes a non-WT amino acid). For genotype quadruplet (G1=00000, G2=00100, G3=00001, G4=00101) with fitness values f1, f2, f3, f4, we calculated |Δf1|=|f2 - f1| and |Δf2|=|f4 - f3|. Buffering of fitness effects by a non-WT amino acid state was quantified as the skewness of the distribution of |Δf1|-|Δf2|, which captures the tendency to reduce the magnitude of fitness effects of substitutions.

### DMS Library Construction

To experimentally validate the predictions that super compensators can increase mutational robustness, one representative super compensator was selected, that is S189A. The sites for random mutagenesis were selected randomly, which are positions 181 and 202.

#### Plasmid Construction

To construct the libraries, we first identified naturally occurring, or, when necessary, inserted restriction sites that were in close proximity to the desired super compensators sites and mutagenesis sites within the *S. cerevisiae HIS3* gene. Single-stranded oligonucleotides that coded for the desired amino acid substitutions (including all synonymous nucleotide variants) were synthesized with 15-25 bp homology arms flanking restriction sites.

Genomic DNA was extracted (Multi Sciences, MK010001) from the prototrophic S288C derived FY4 strain. The *HIS3* locus (ATG – 250 bp, TAG + 300 bp) was then amplified using 500 ng of the extracted DNA, primers 376, 379, and the Golden Star T6 DNA polymerase (Tsingke Biotech, TSE101). The amplicon was PCR purified (Qiagen MinElute, 28004) and inserted between the SacI and KpnI sites (destroying them) within the pGREG506 ADH1pr-MCM-mCherry vectors (a gift from the Chao Tang’s lab) via Gibson Assembly (Tiangen Easygeno, VI201) to yield plasmid p142. After PCR, both amplicons were verified, purified, and assembled into the vector backbone to yield plasmid p143.

### Library Construction

To construct the final libraries, 7.5 μg of p142 and p143 were digested for 4 hours at 37°C with 9 μL of KpnI, NheI (NEB, R3131S), and BstXI (NEB, R01131S) and EagI (NEB, R3505S) respectively. Each reaction was then loaded on a 0.7% agarose gel, with 0.5X TAE running buffer, gel purified (Magen, D2111), and concentrated (Tianmo Biotech, TD413). Subsequently, the library ssOligos (primers 337, 338) were then assembled into 500 ng of the digested vectors to a final concentration of 0.25 μM in 15 μL Gibson Assembly reactions (2 step Gibson Assembly, 3 minutes at 37°C, 60 minutes at 50°C). Finally, the assembly reactions were split into 5 transformations in chemically competent *E. Coli* (Transgen Biotech, CD101) and spread onto LB+Amp plates, and incubated overnight (∼14 hours) at 37°C. After incubation, the colonies were verified (colony PCR, Sanger sequencing) and scraped off the plates (∼55,000 colonies/library), mixed and the library plasmids were extracted (Tiangen Biotech, DP106), quantified, and stored.

### Yeast Transformation and Bulk Competition

Competent yeast cells were prepared (Zymo Research, T2001) using yeast strain Y120 (derived from UCC8363; MKT1-RM SAL1-RM CAT5-RM MIP1-BY ura3Δ0 *his3*:KanMX). Subsequently, 5 μg of each plasmid was transformed into ∼52.5 x 10^6^ cells and incubated at 30°C, 220 RPM for 50 hrs in 1L synthetic complete dropout medium (SCD) lacking Uracil (-Ura).

After transformation (yielding ∼750,000 colonies/library), ∼3 x 10^8^ cells per library were inoculated into 1 L of SCD-Ura and grown at 30°C, 220 RPM for 6 hrs. After that, ∼3 x 10^7^ cells per library were inoculated into 1 L SCD-His (Histidine) and ∼2.1 x 10^7^ cells into 700 mL SCD-Ura and incubated at 30°C, 220 RPM. The competition was carried out for 120 hours with 12 hours between bottlenecks (2.1 x 10^7^ cells per library transferred into fresh media), two samples of 15 mL (technical replicates) were taken from each culture every 24 hours, centrifuged (3000 g, 5 mins) and stored (-20°C), the experiment from transformation to completion was carried out twice (biological replicates).

### Plasmid Extraction and Illumina Sequencing

After competition, cell pellets from each time point were thawed and plasmids extracted using the Tiangen Midiprep kit (DP106). Extracted DNA (∼10 ng) was labeled and amplified using staggered primers (primers 345-356) and Q5 DNA polymerase (NEB, M0492S) in 50 μL reactions (98°C for the 30 s; 98°C for 10 s, 55°C for 30 s and 72°C for 30 s for 17 cycles; 72°C for 2 min). The amplicons were purified (Qiagen MinElute, 28004) and pooled samples were submitted to Genewiz company for Illumina library construction and sequencing.

### Data Processing

Raw paired-end FASTQ files were merged using FLASH (Magoč and Salzberg 2011), demultiplexed according to the unique primer pairs used to amplify each sample at specific timepoint, and reads for each genotype were counted using starcode (Zorita et al. 2015). Reads for each genotype were the sum of all synonymous nucleotide genotypes and the sum across two technical replicates. Fitness for each genotype was calculated using PyFitSeq (Li et al. 2018), and the average fitness across both biological replicates was used for further analysis.

### Prevalence Analysis of Super Compensators

A total of 217 DMS datasets covering 186 unique proteins were retrieved from the ProteinGym database. These datasets incorporate quantitative fitness measurements derived from diverse biochemical and cellular readouts, including protein stability, enzymatic activity, and bacterial growth rate. To detect potential super-compensators and quantify their buffering capacity against substitutions, the established computational pipeline was applied across these datasets. The filtering criteria required datasets to contain multiple substitutions per genotype—specifically at least 10 sets of quadruplet genotypes—to evaluate whether a given amino acid state could mitigate mutational effects. This filtering process yielded four eligible datasets: three assayed for in vitro protein folding stability using cDNA display proteolysis: E3 ubiquitin-protein ligase AMFR, iron-regulated surface determinant protein H, and E3 SUMO-protein ligase CBX4 (Tsuboyama et al., 2023), and one assayed for fluorescence intensity via FACS (green fluorescent protein 2) (Gonzalez Somermeyer et al., 2022).

## Data, Materials, and Software Availability

All data and codes used for analysis and creating figures are available on Zenodo at https://doi.org/10.5281/zenodo.17769277

## Acknowledgments

We thank Dr. Chu Wang and Dr. Luhua Lai at Peking University for helpful discussions and comments on the manuscript. This work is funded by China Postdoctoral Science Foundation (2025M782617 to W.H.J.), the Natural Science Foundation of Hubei Province (JCZRQNB202600450 to W.H.J), Open Project Funding of the Key Laboratory of Fermentation Engineering (Ministry of Education) (202509FE11 to W.H.J.), the Natural Science Foundation of Xinjiang Uygur Autonomous Region (2025D01A136, XJRC-2025-KJ-YJCXPT-103 to H.C.Y.). Additional funds to L.B.C. are provided by Peking-Tsinghua Center for Life Sciences.

## Author Contributions

L.B.C. and W.H.J. designed the study; W.H.J., S.S.K., H.C.Y., Yiparemu and L.B.C. carried out the experiments and analyzed data; W.H.J., H.C.Y., S.S.K. and L.B.C wrote the manuscript. All authors read and approved the final version of the manuscript.

## Supplementary Figures and Tables

**Fig. S1.**
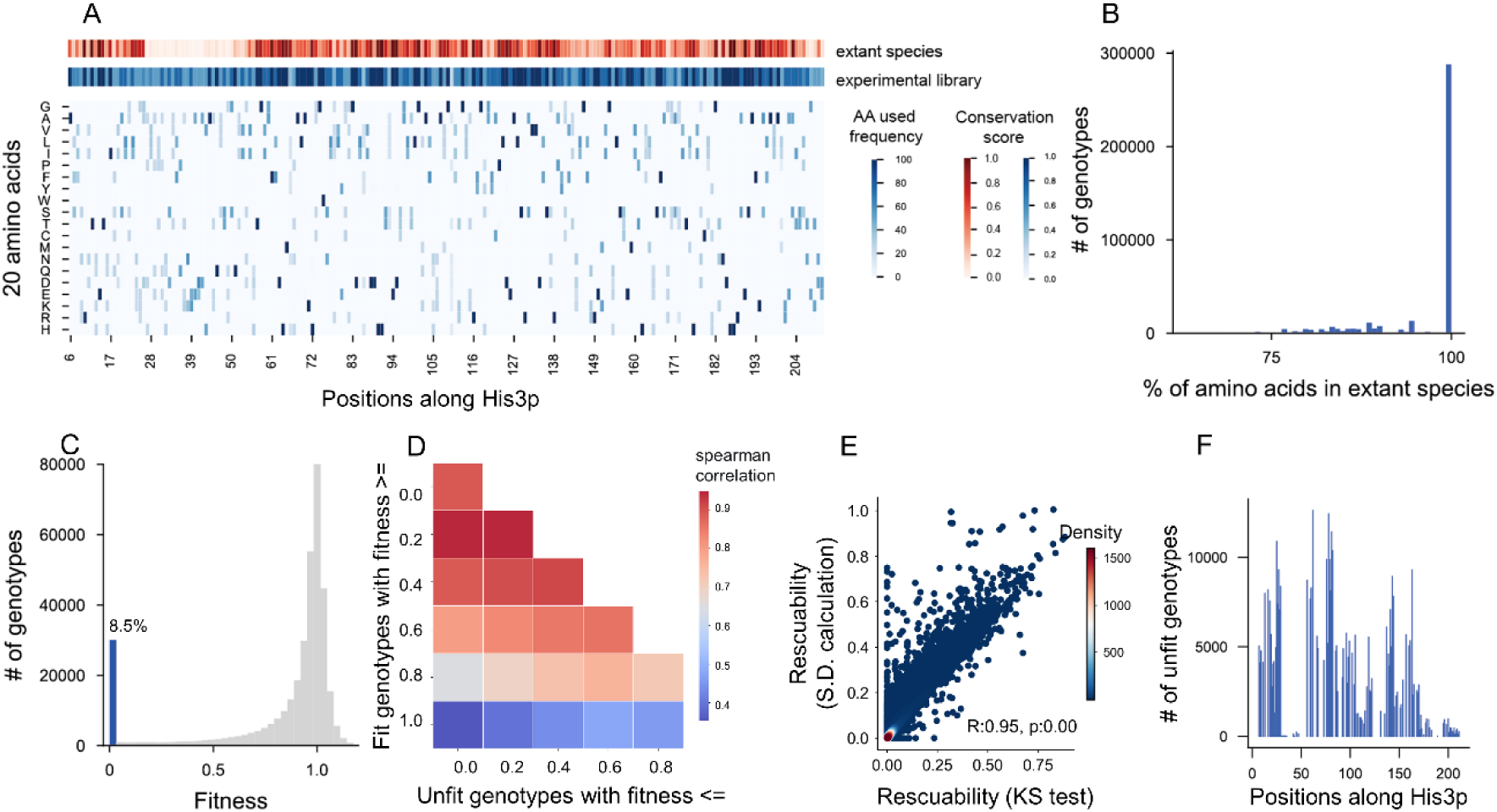
The DMS library is derived from extant yeast sequences and spans nearly the full length of His3p. (A) Position wise conservation and amino acid conservation across His3p. The red track shows conservation score across 21 extant yeast species at each position of the His3p monomer. The blue track shows the conservation score in the DMS library with the heatmap below showing the frequency of each amino acid in each position in the library. (B) Most genotypes in the library are composed of extant amino acids. (C) Fitness distribution of all genotypes in the library, the blue bar indicates percentage of unfit genotypes. (D) We defined “rescue” as a fitness increase from 0 to a value significantly above 0, and calculated rescuability accordingly. The correlation between rescuability values was then examined across a range of fitness thresholds used to define unfit and fit genotypes. (E) Different rescuability calculation methods produce strong agreement of rescuability estimates for unfit genotypes (R_2_ = 0.84). (F) Unfit genotypes span nearly the entire length of His3p.

**Fig. S2.**
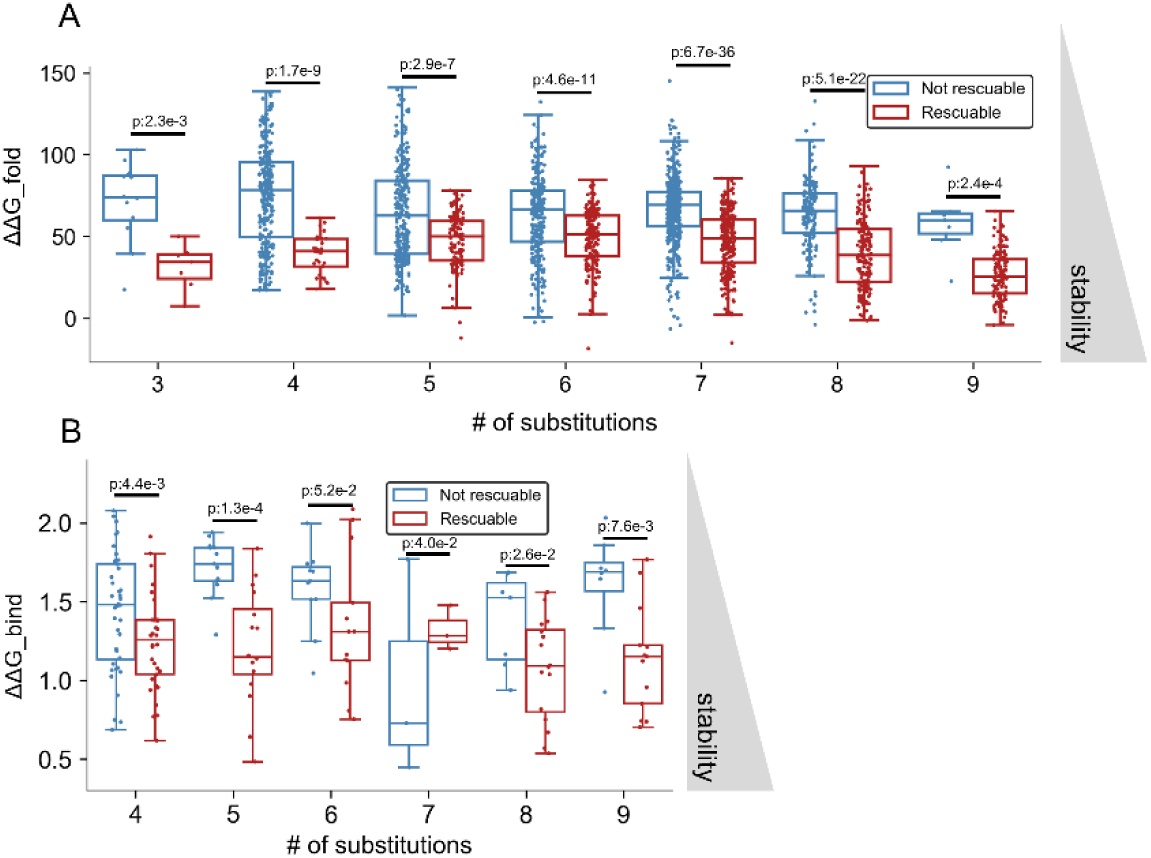
Rescuable genotypes are more structurally stable than non-rescuable genotypes. (A) Across varying numbers of substitutions, rescuable genotypes display higher predicted structural stability than non-rescuable genotypes. (B) Among genotypes with more than 50% substitutions in subunit interface regions, rescuable genotypes remain more structurally stable than non-rescuable genotypes across varying numbers of substitutions.

**Fig. S3.**
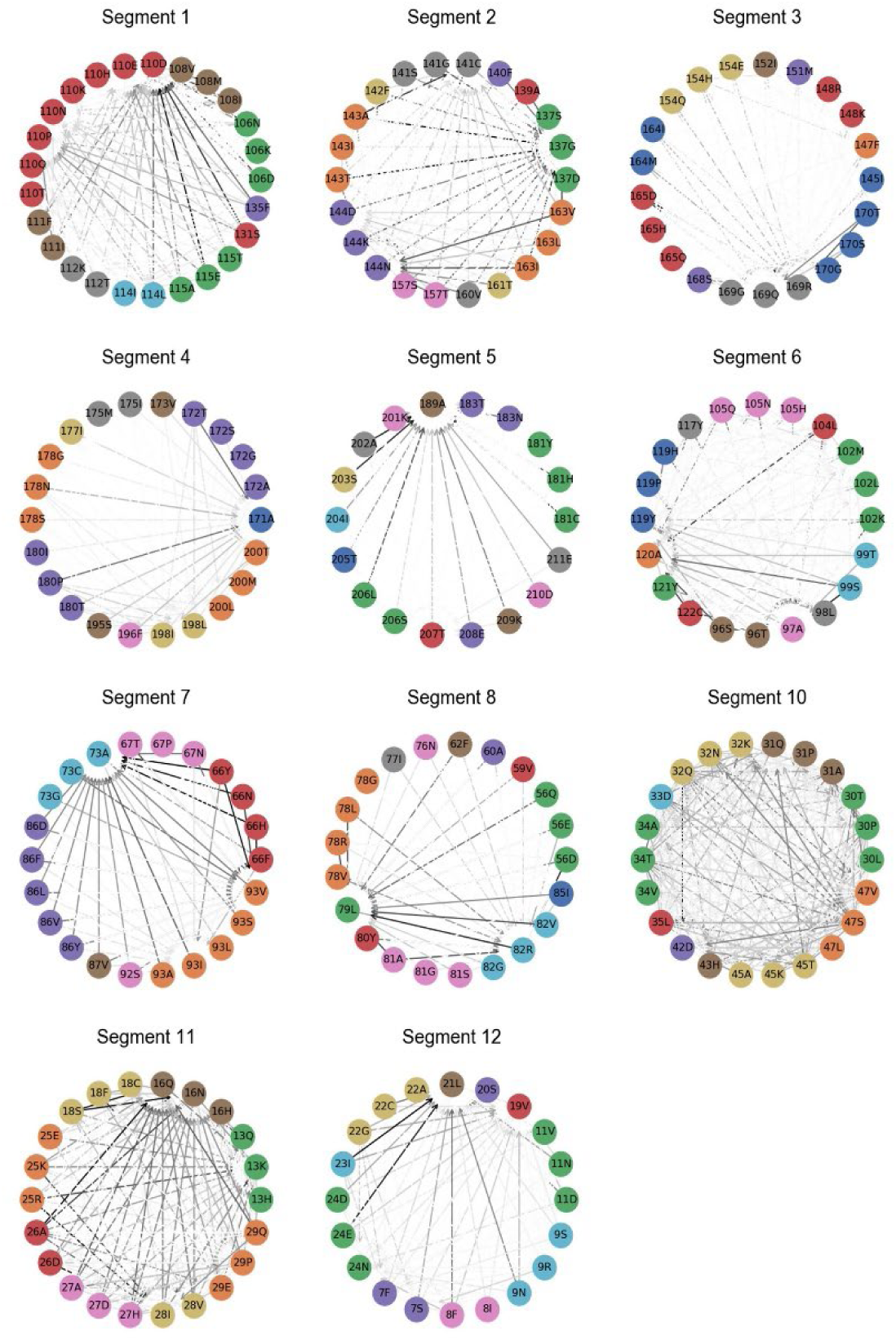
There exist several substitutions that increase fitness of diverse genotypes. Compensatory relationships of all pairs of substitutions and non-WT amino acid states at distinct positions for all segments, excluding segment 9.

**Fig. S4.**
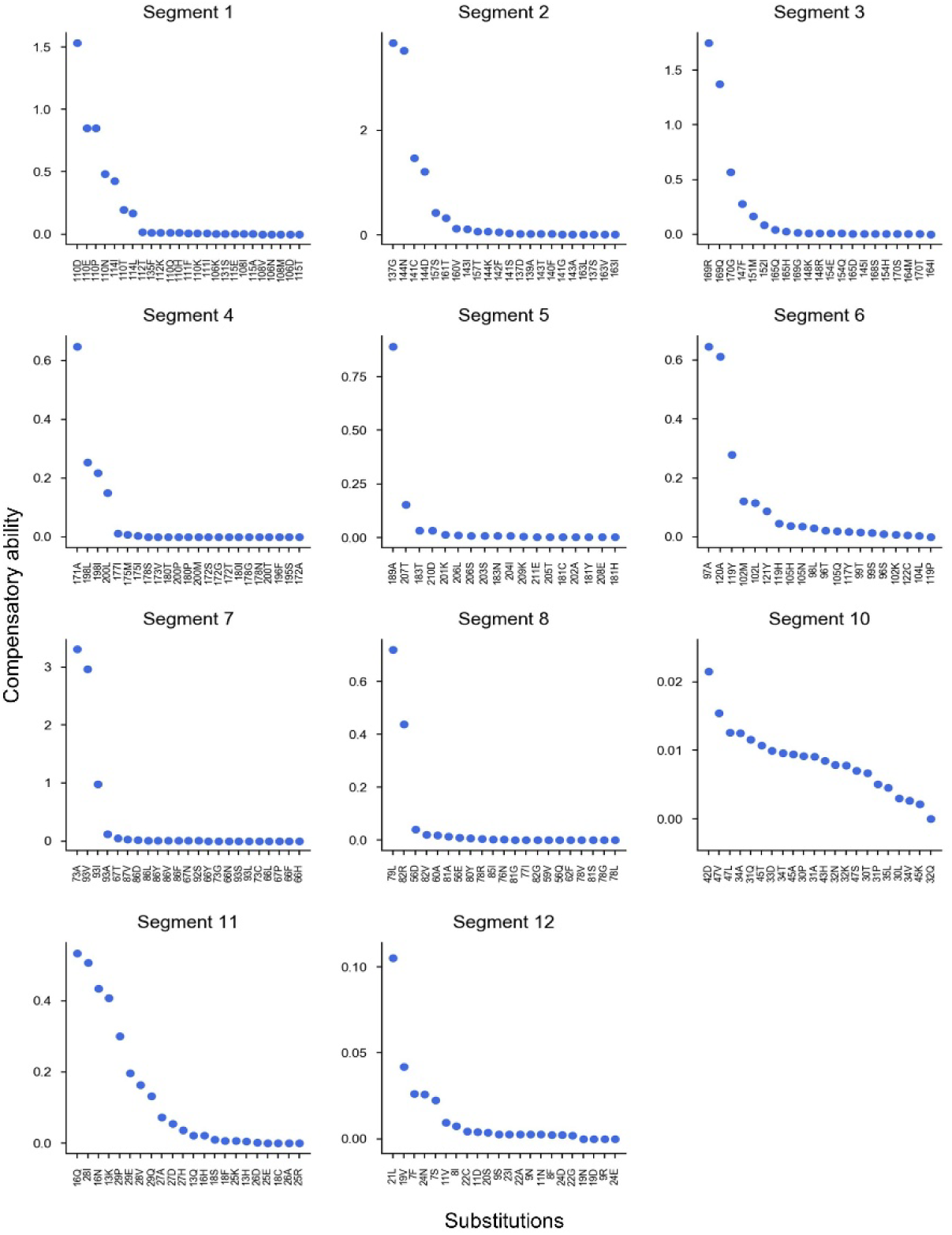
Several substitutions can increase fitness across diverse genetic backgrounds. Compensatory capacity of each substitution across all segments, excluding segment 9.

**Fig. S5.**
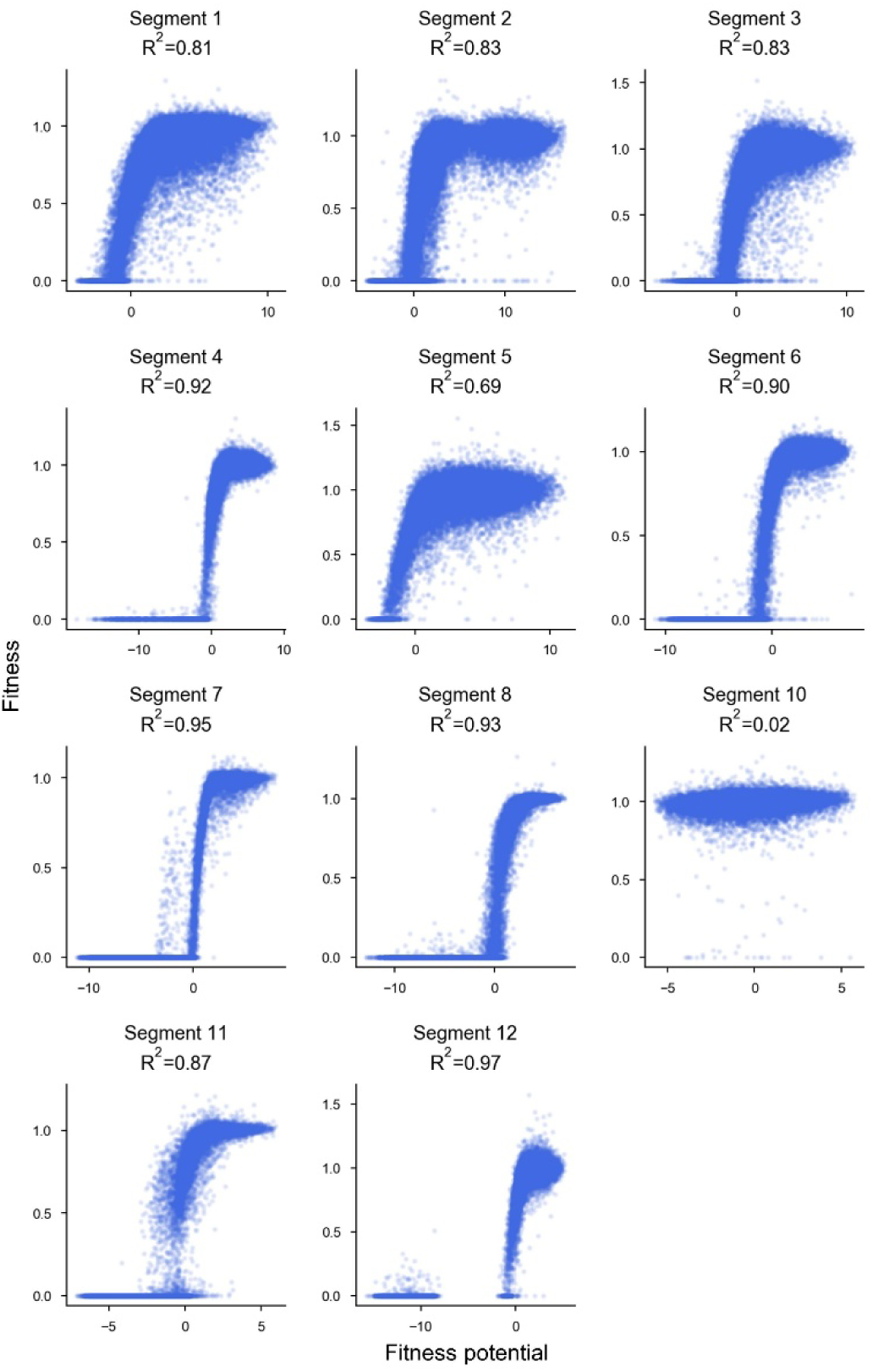
Fitness potential is a latent variable that can predict fitness. Each panel illustrates the non-linear mapping learned by a neural network model for each of the 12 His3p segments analyzed, excluding segment 9.

**Fig. S6.**
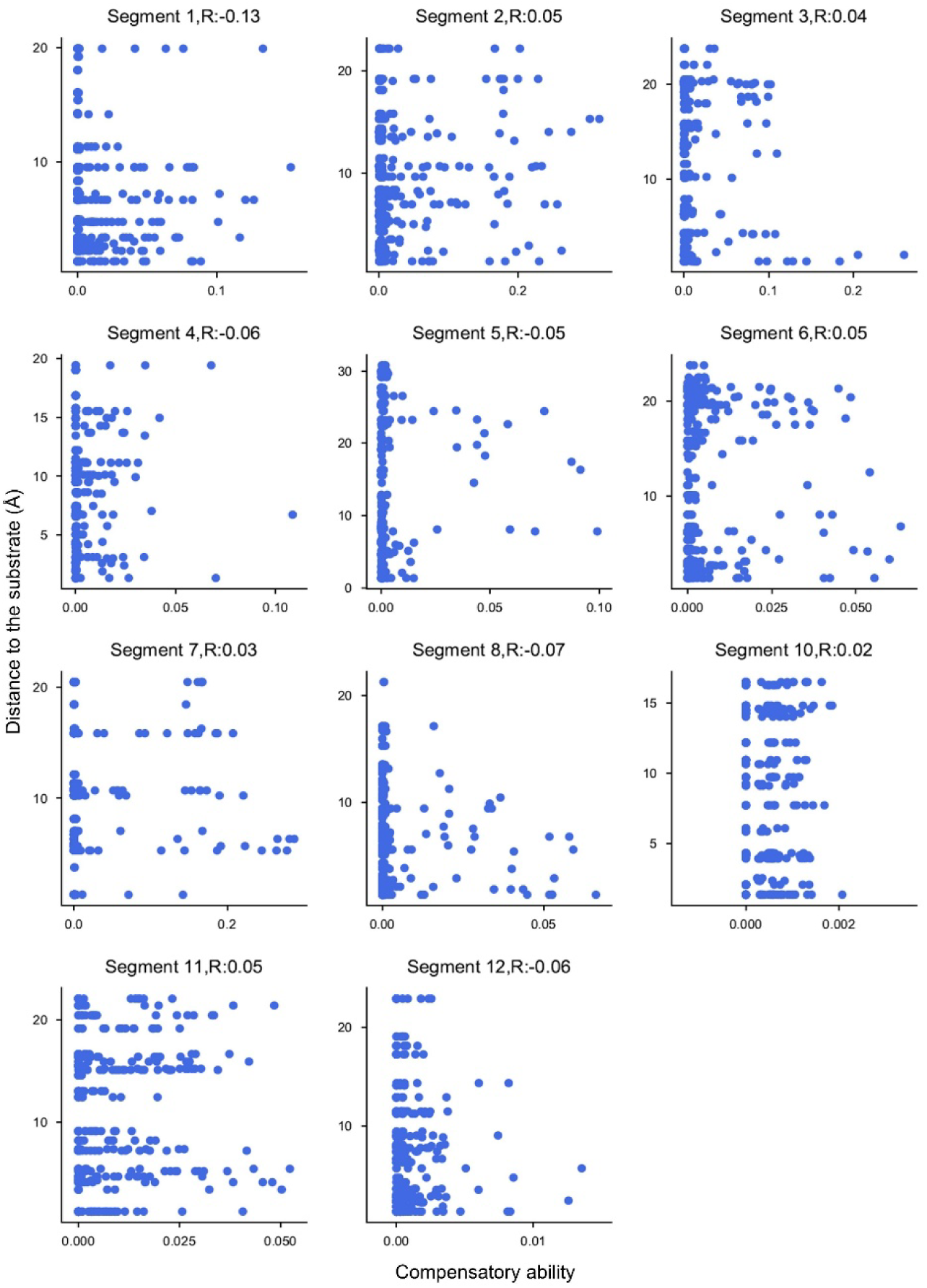
Distance from substituted position of each non-WT amino acid state to the substrate. Distance from each non-WT amino acid state to the substrate (Å, y-axis) was plotted against compensatory ability (x-axis) for all segments except segment 9. There is no significant correlation between the distance from each non-WT amino acid state to the substrate and its compensatory ability.

**Fig. S7.**
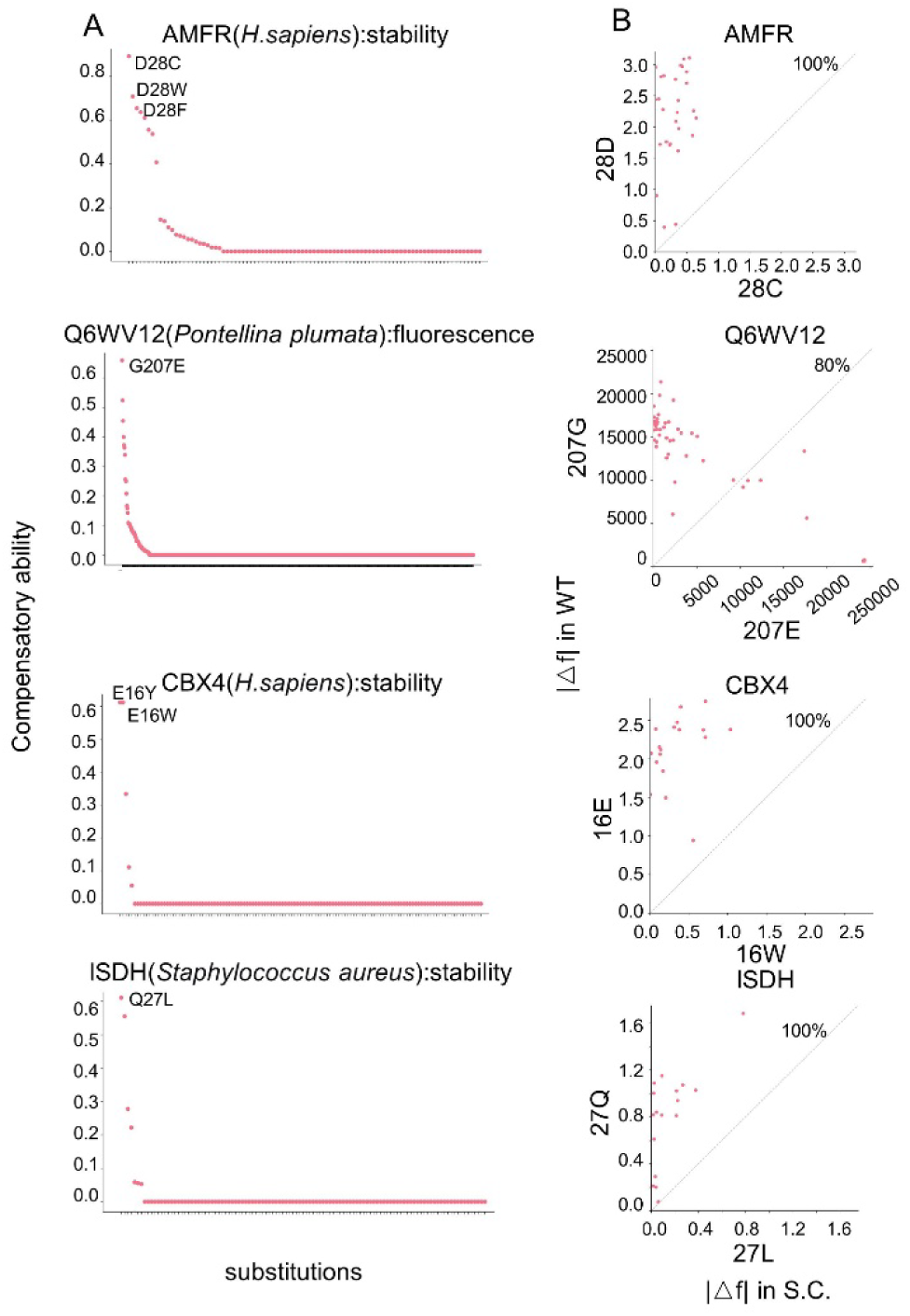
Super compensators identified across diverse proteins. (A) Distribution of compensatory ability for individual substitution across different proteins. Substitutions with prominent compensatory ability (≥ 0.6) are labeled. The fitness metric used in each dataset (stability or fluorescence intensity) is indicated in the title of each panel. (B) Super compensators buffer the fitness effects of other substitutions. Each panel corresponds to the super compensator highlighted in panel A (which showed the highest compensatory ability for that protein). The percentage of substitutions exhibiting reduced fitness effects in the super compensator background is labeled.

**Fig. S8.**
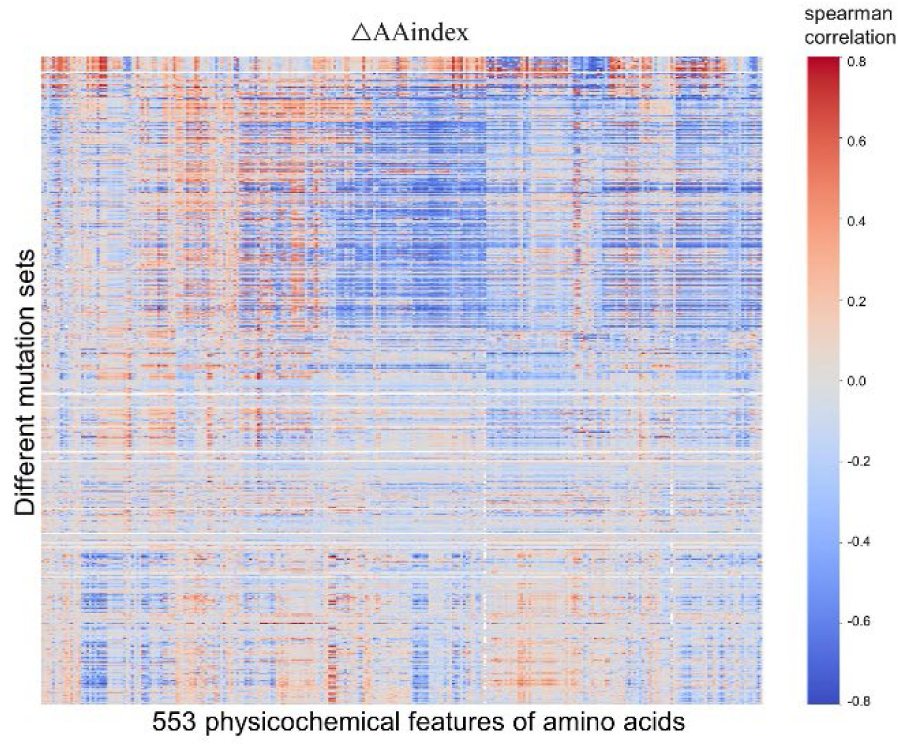
Amino acid physicochemical properties can predict fitness. The heatmap shows Spearman correlations between fitness and physicochemical properties (AAindex values) distance from WT. Each column corresponds to one of 553 AAindex descriptors, and each row corresponds to a mutation set (genotypes carrying substitutions at the same positions). For each mutation set and AAindex descriptor, we computed the Spearman correlation (ρ) between the AAindex distance to the wild-type sequence and fitness across all genotypes in that mutation set. P values were adjusted for multiple testing using the Benjamini–Hochberg false discovery rate (FDR) procedure, and non-significant correlations (FDR ≥ 0.05) were set to zero. Columns (AAindex descriptors) were hierarchically clustered using Ward’s linkage on the matrix of correlation coefficients (top dendrogram). Rows (mutation sets) were ordered by the fraction of substituted residues located in structural loop regions (left grayscale bar, 0–100%). The heatmap shows the correlation coefficients (blue, negative; red, positive; white, ρ ≈ 0). The x-axis label indicates the total number of AAindex descriptors, and the y-axis label denotes mutation sets sorted by loop fraction.

**Fig. S9.**
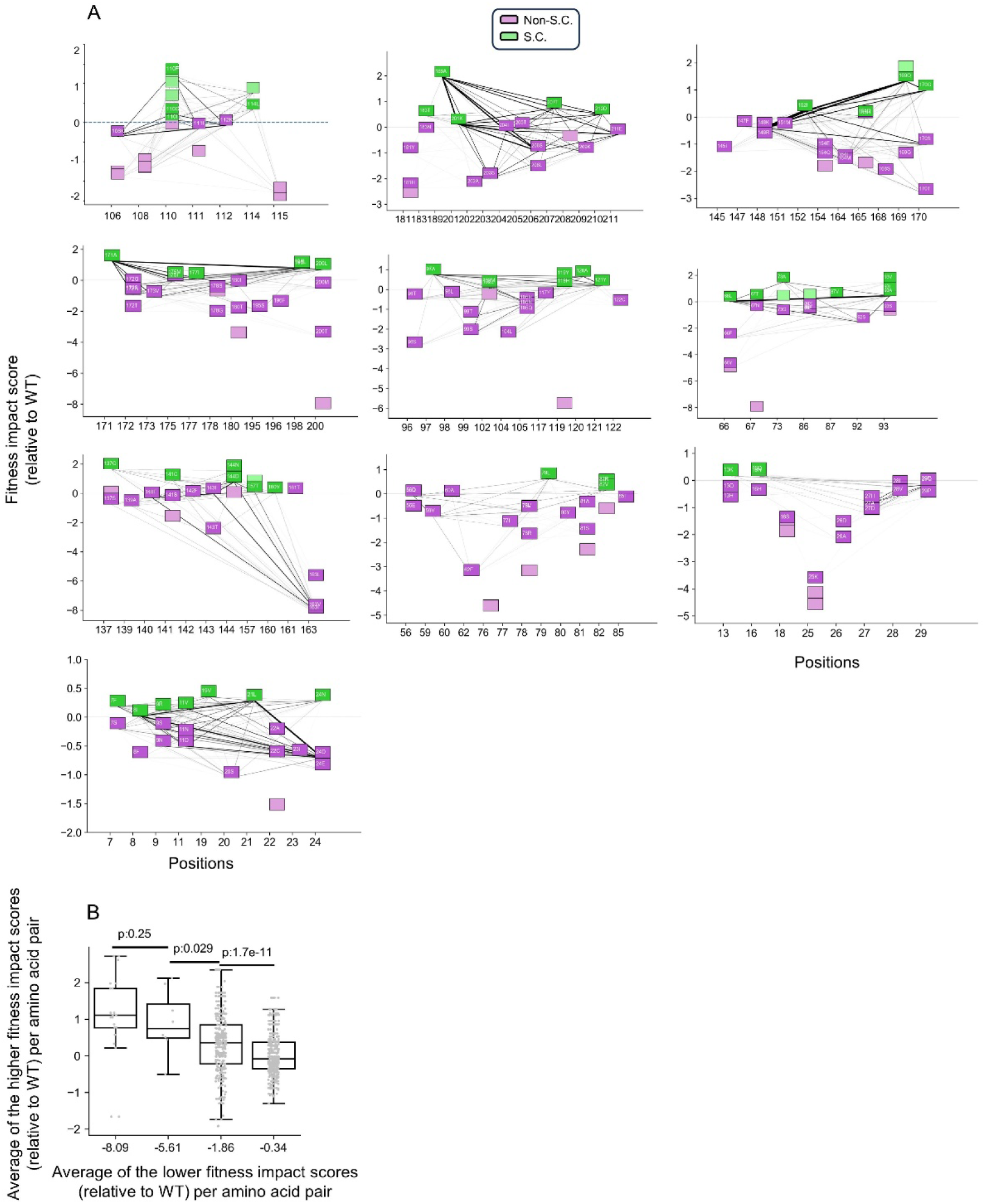
Co-occurring amino-acid state pairs in extant His3p orthologs, across all segments excluding segments 9 and 10. (A) Co-occurrence network of amino acid state pairs across 355 extant yeast species. Nodes represent amino acid states and edges the co-occurrence of a pair. Edge shading reflects the fraction of species in which a given pair co-occurs. Non-extant amino acid states are denoted by lighter node shading. Pairs that co-occur in at least 10 species are labeled with white text. (B) Among pairs of non-WT amino-acid states observed in extant species, amino-acids states with lower fitness impact scores relative to WT (x-axis, binned) tend to co-occur with states exhibiting higher fitness-impact scores (y-axis).

**Table S1.**
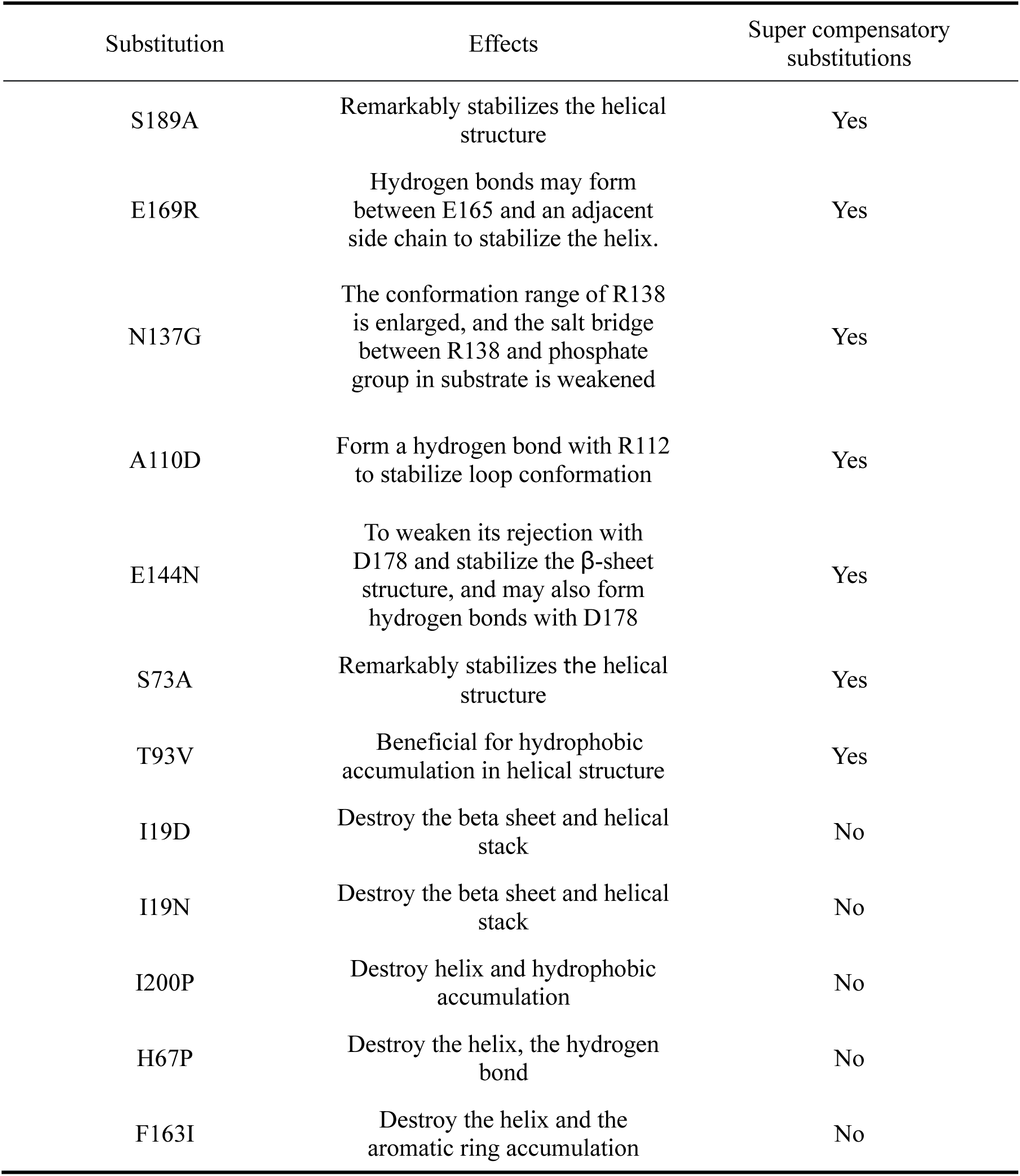
Structural effects of super compensatory and non-compensatory substitutions. Predicted structural effects of 7 super compensators (highest fitness impact score relative to WT) and five non compensatory substitutions (lowest fitness impact score relative to WT).

