## Supplemental Files for "Super compensatory substitutions restore protein function and confer mutational robustness"

### Supplementary Figures and Tables

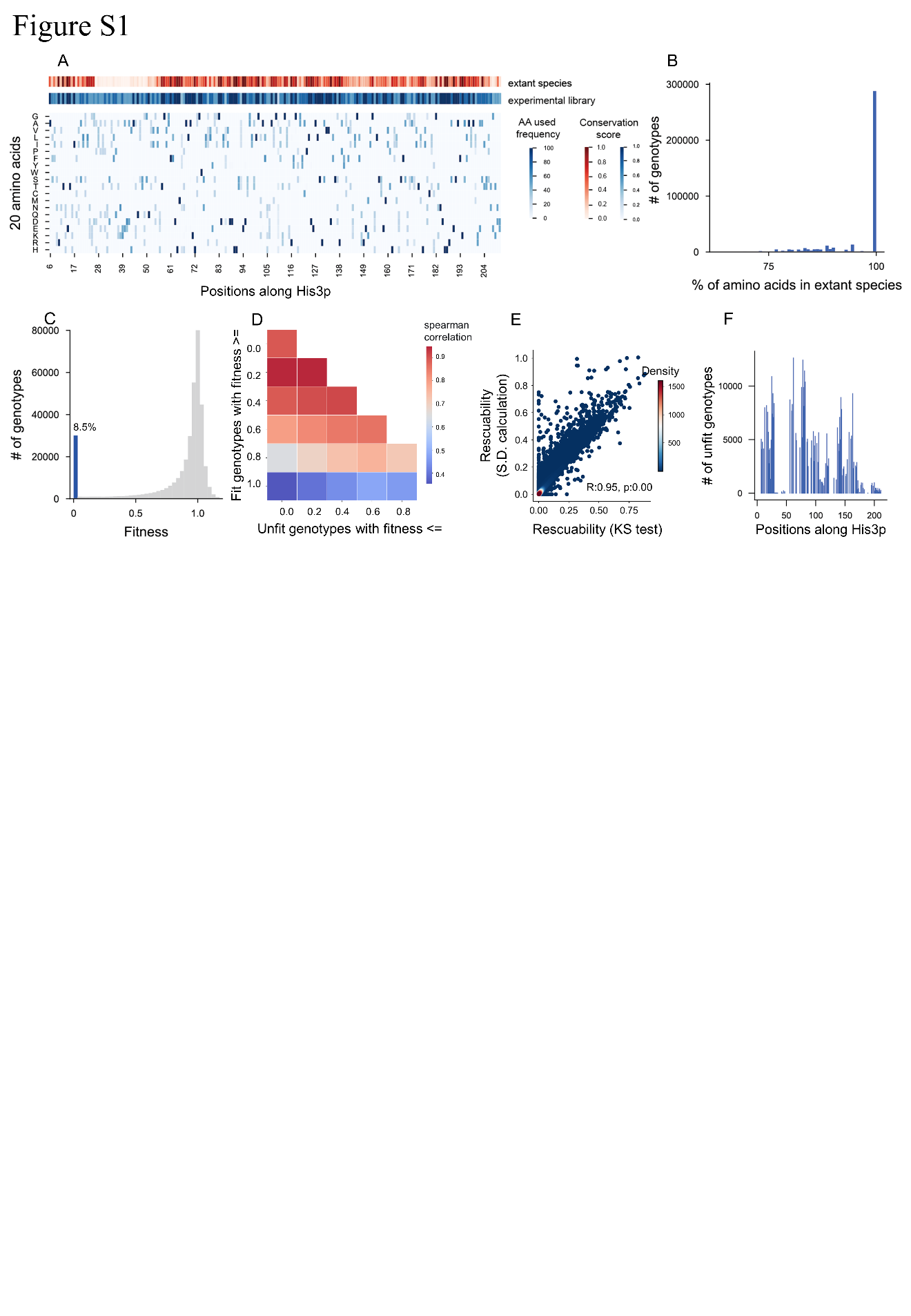
**Fig. S1. The DMS library is derived from extant yeast sequences and spans nearly the full length of His3p.** (A) Position wise conservation and amino acid conservation across His3p. The red track shows conservation score across 21 extant yeast species at each position of the His3p monomer. The blue track shows the conservation score in the DMS library with the heatmap below showing the frequency of each amino acid in each position in the library. (B) Most genotypes in the library are composed of extant amino acids. (C) Fitness distribution of all genotypes in the library, the blue bar indicates percentage of unfit genotypes. (D) We defined “rescue” as a fitness increase from 0 to a value significantly above 0, and calculated rescuability accordingly. The correlation between rescuability values was then examined across a range of fitness thresholds used to define unfit and fit genotypes. (E) Different rescuability calculation methods produce strong agreement of rescuability estimates for unfit genotypes (R^2^ = 0.84). (F) Unfit genotypes span nearly the entire length of His3p.

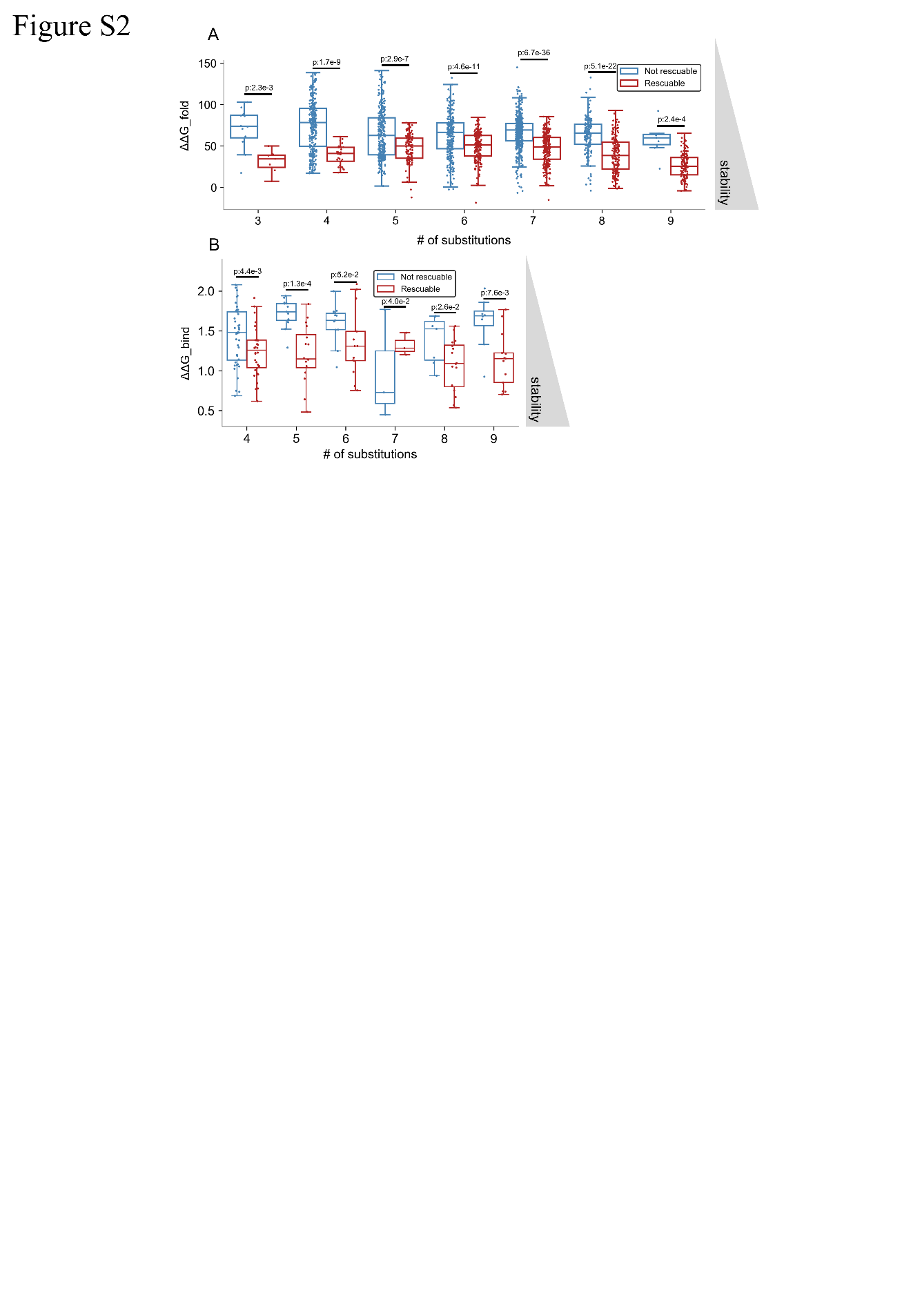
**Fig. S2. Rescuable genotypes are more structurally stable than non-rescuable genotypes.** (A) Across varying numbers of substitutions, rescuable genotypes display higher predicted structural stability than non-rescuable genotypes. (B) Among genotypes with more than 50% substitutions in subunit interface regions, rescuable genotypes remain more structurally stable than non-rescuable genotypes across varying numbers of substitutions.

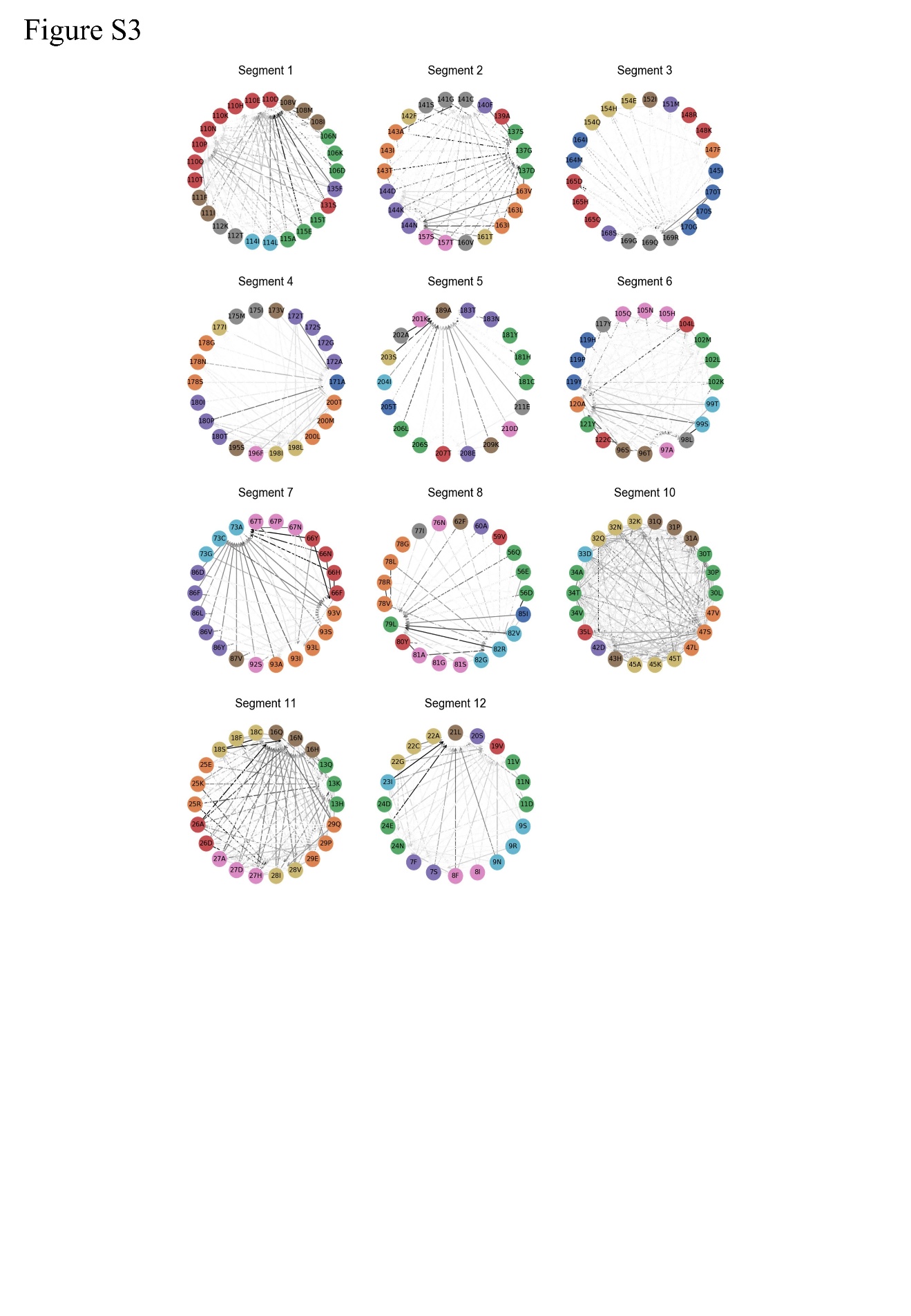
**Fig. S3.** **There exist several substitutions that increase fitness of diverse genotypes.** Compensatory relationships of all pairs of substitutions and non-WT amino acid states at distinct positions for all segments, excluding segment 9.

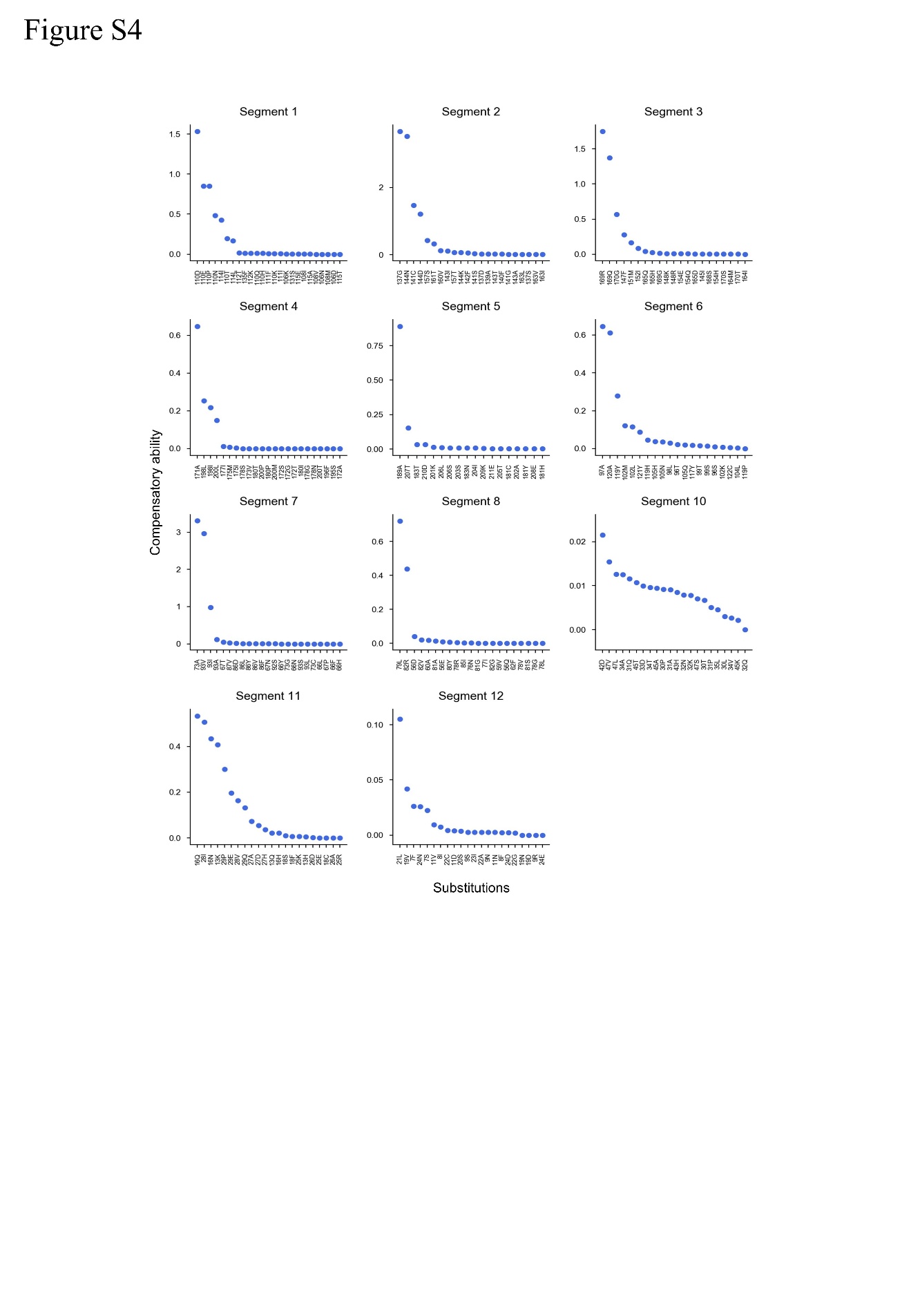

**Fig. S4.** **Several substitutions can increase fitness across diverse genetic backgrounds.** Compensatory capacity of each substitution across all segments, excluding segment 9.

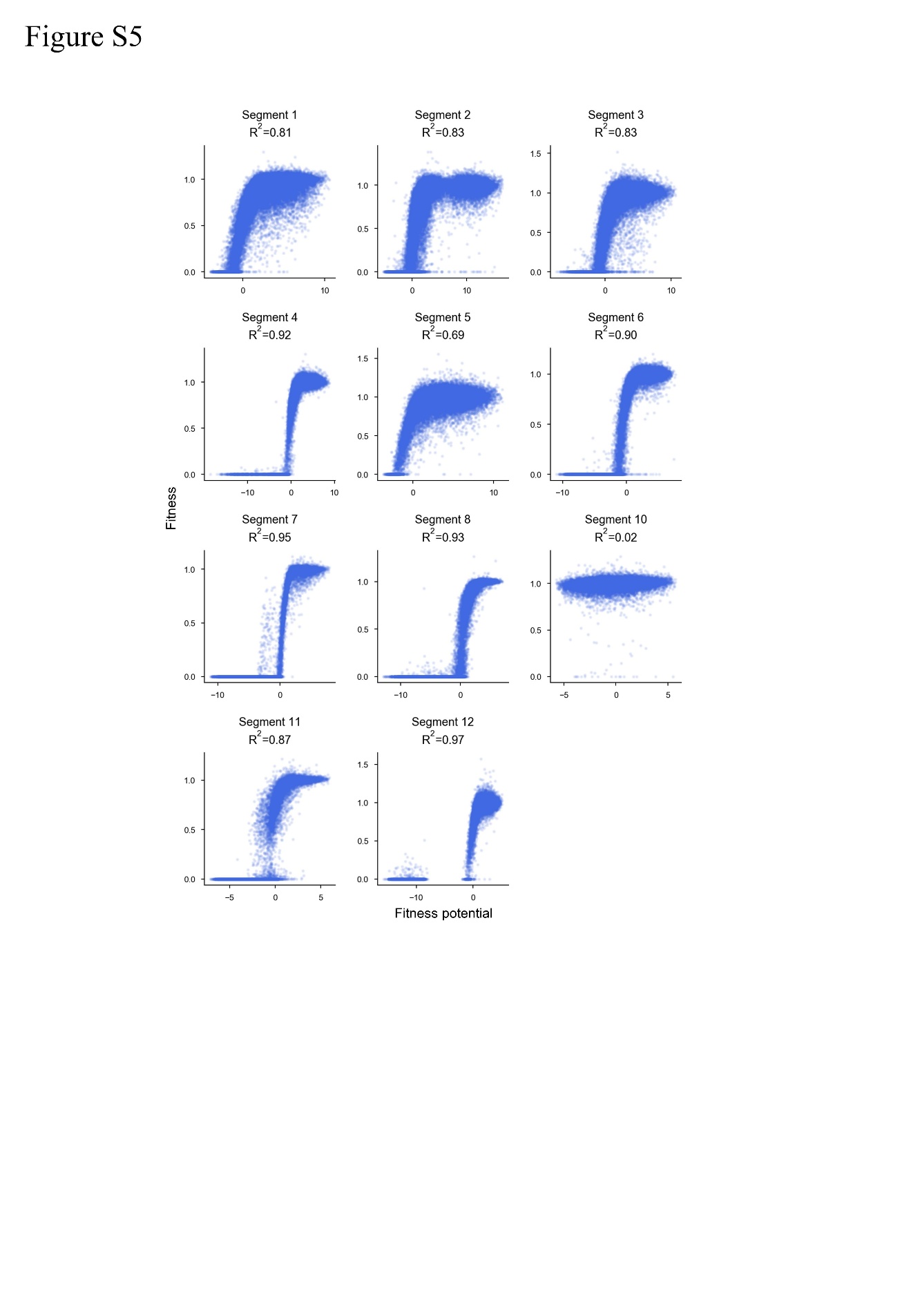

**Fig. S5. Fitness potential is a latent variable that can predict fitness.** Each panel illustrates the non-linear mapping learned by a neural network model for each of the 12 His3p segments analyzed, excluding segment 9.

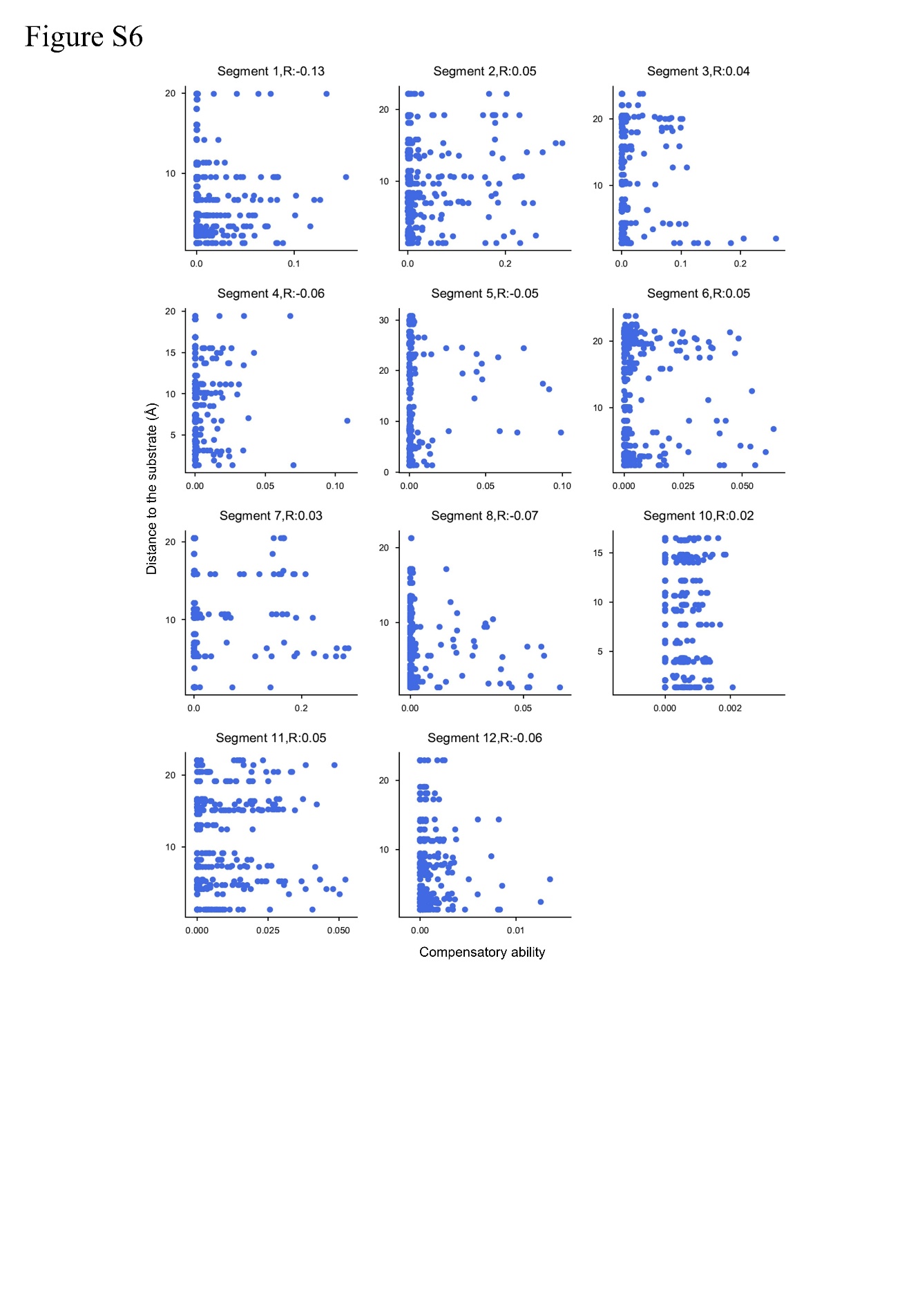

**Fig. S6.** **Distance from substituted position of each non-WT amino acid state to the substrate.** Distance from each non-WT amino acid state to the substrate (Å, y-axis) was plotted against compensatory ability (x-axis) for all segments except segment 9. There is no significant correlation between the distance from each non-WT amino acid state to the substrate and its compensatory ability.

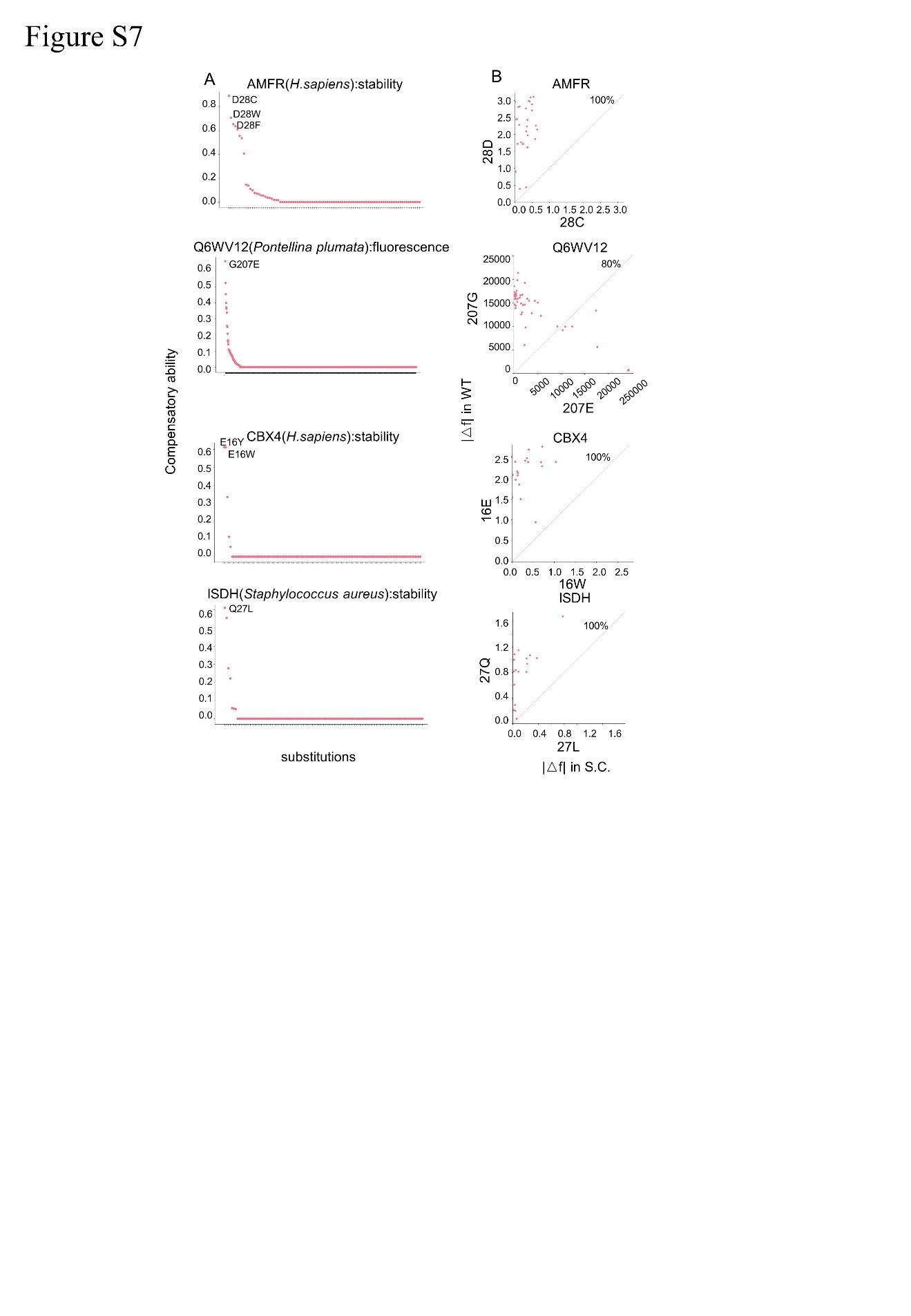

**Fig. S7. Super compensators identified across diverse proteins.** (A) Distribution of compensatory ability for individual substitution across different proteins. Substitutions with prominent compensatory ability ( ≥ 0.6) are labeled. The fitness metric used in each dataset (stability or fluorescence intensity) is indicated in the title of each panel. (B) Super compensators buffer the fitness effects of other substitutions. Each panel corresponds to the super compensator highlighted in panel A (which showed the highest compensatory ability for that protein). The percentage of substitutions exhibiting reduced fitness effects in the super compensator background is labeled.

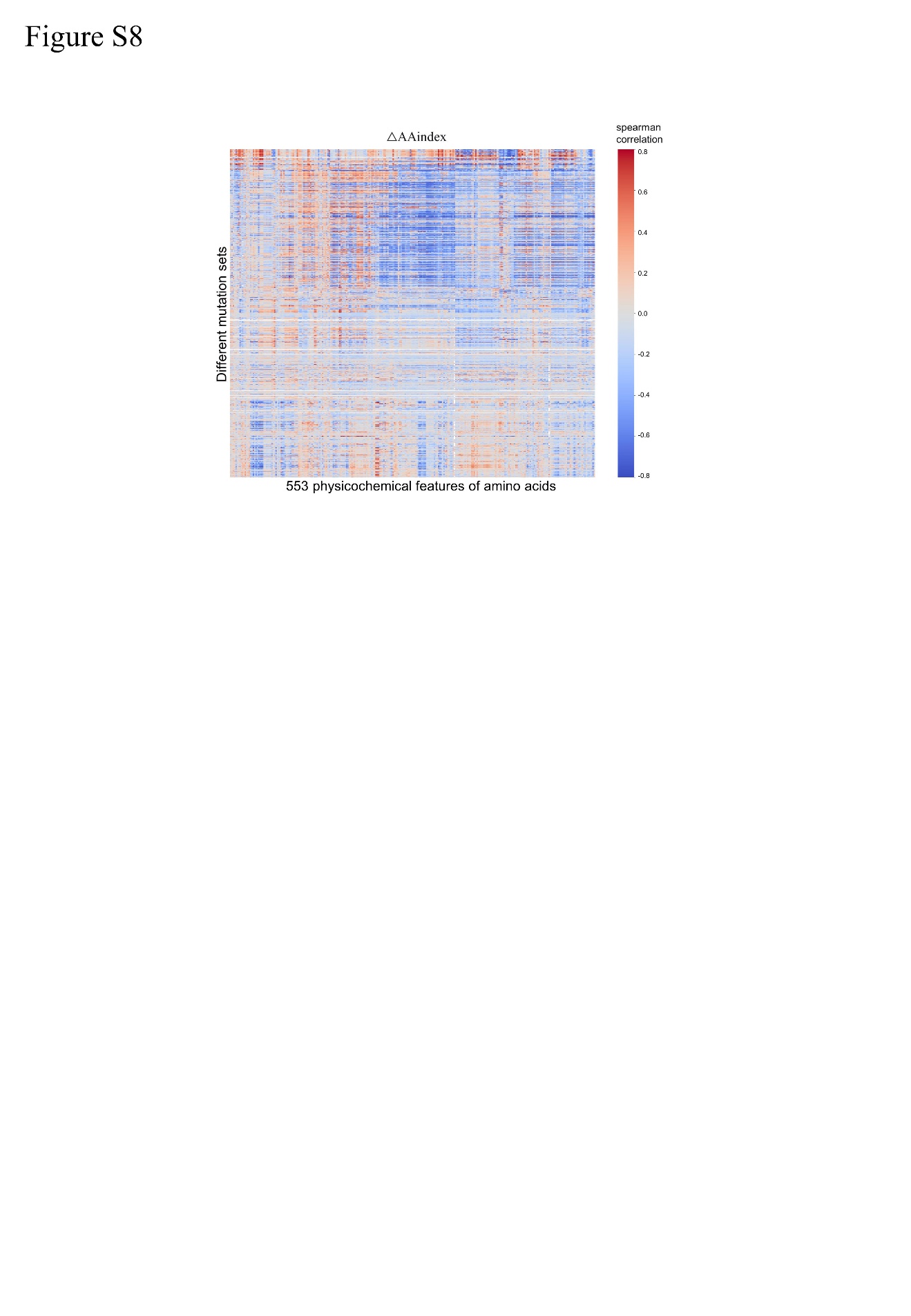

**Fig. S8. Amino acid physicochemical properties can predict fitness.** The heatmap shows Spearman correlations between fitness and physicochemical properties (AAindex values) distance from WT. Each column corresponds to one of 553 AAindex descriptors, and each row corresponds to a mutation set (genotypes carrying substitutions at the same positions). For each mutation set and AAindex descriptor, we computed the Spearman correlation (ρ) between the AAindex distance to the wild-type sequence and fitness across all genotypes in that mutation set. P values were adjusted for multiple testing using the Benjamini–Hochberg false discovery rate (FDR) procedure, and non-significant correlations (FDR ≥ 0.05) were set to zero. Columns (AAindex descriptors) were hierarchically clustered using Ward’s linkage on the matrix of correlation coefficients (top dendrogram). Rows (mutation sets) were ordered by the fraction of substituted residues located in structural loop regions (left grayscale bar, 0–100%). The heatmap shows the correlation coefficients (blue, negative; red, positive; white, ρ ≈ 0). The x-axis label indicates the total number of AAindex descriptors, and the y-axis label denotes mutation sets sorted by loop fraction.

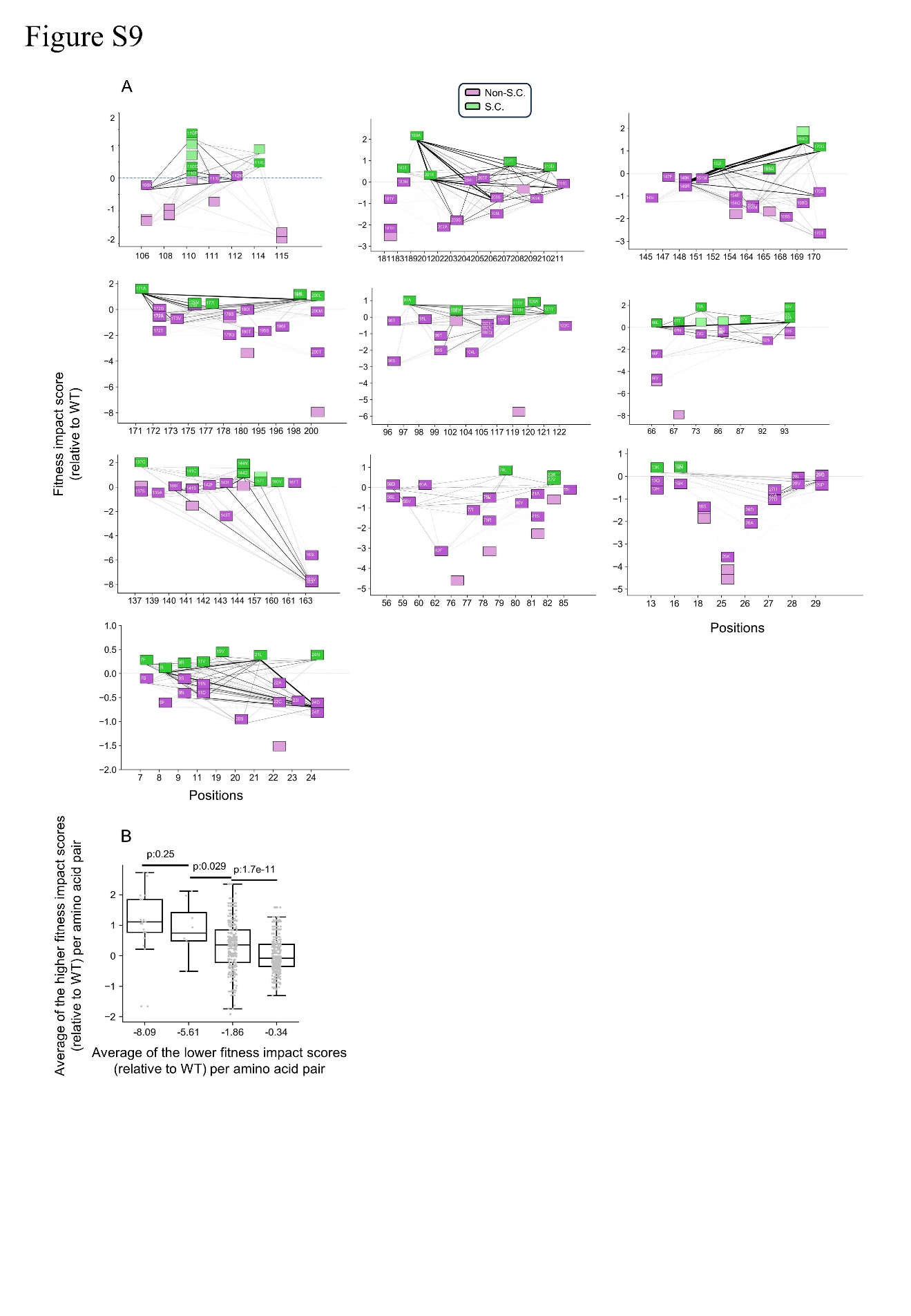

**Fig. S9. Co‑occurring amino‑acid state pairs in extant His3p orthologs, across all segments excluding segments 9 and 10.** (A) Co-occurrence network of amino acid state pairs across 355 extant yeast species. Nodes represent amino acid states and edges the co-occurrence of a pair. Edge shading reflects the fraction of species in which a given pair co-occurs. Non-extant amino acid states are denoted by lighter node shading. Pairs that co-occur in at least 10 species are labeled with white text. (B) Among pairs of non‑WT amino‑acid states observed in extant species, amino-acids states with lower fitness impact scores relative to WT (x-axis, binned) tend to co-occur with states exhibiting higher fitness-impact scores (y-axis).

**Table S1. Structural effects of super compensatory and non-compensatory substitutions.** Predicted structural effects of 7 super compensators (highest fitness impact score relative to WT) and five non compensatory substitutions (lowest fitness impact score relative to WT)**.**

| Substitution | Effects | Super compensatory substitutions |
| --- | --- | --- |
| S189A | Remarkably stabilizes the helical structure | Yes |
| E169R | Hydrogen bonds may form between E165 and an adjacent side chain to stabilize the helix. | Yes |
| N137G | The conformation range of R138 is enlarged, and the salt bridge between R138 and phosphate group in substrate is weakened | Yes |
| A110D | Form a hydrogen bond with R112 to stabilize loop conformation | Yes |
| E144N | To weaken its rejection with D178 and stabilize the β-sheet structure, and may also form hydrogen bonds with D178 | Yes |
| S73A | Remarkably stabilizes the helical structure | Yes |
| T93V | Beneficial for hydrophobic accumulation in helical structure | Yes |
| I19D | Destroy the beta sheet and helical stack | No |
| I19N | Destroy the beta sheet and helical stack | No |
| I200P | Destroy helix and hydrophobic accumulation | No |
| H67P | Destroy the helix, the hydrogen bond | No |
| F163I | Destroy the helix and the aromatic ring accumulation | No |
